# Density-dependent plasticity of female reproductive output mediates Allee effects in *Drosophila melanogaster*

**DOI:** 10.64898/2026.08.08.743646

**Authors:** Rutvij Kaustubh Kulkarni, Yashaswini M. B., Revanth Gowda, Vasu Sheeba

**Author notes:** Corresponding author information: Correspondence may be directed to V.S. or R.K.K. via email- V.S., R.K.K. Data availability: All data used in this manuscript along with the MATLAB code used for custom analyses are available upon request. Author contributions: R.K.K. conceived the project, designed experiments, analysed data, and wrote the manuscript. R.K.K., Y.M.B. and R.G. performed experiments. V.S. supervised the project and wrote the manuscript.

## Abstract

Allee effects are positive relationships between components of individual fitness and the number or density of individuals in a population. Negative density effects are well documented in *Drosophila melanogaster* across life stages, while Allee effects are rare and largely confined to the larval stages. However, despite the costs of high density, adult flies are found to exhibit attraction to same-sex conspecifics, suggesting some fitness value to the presence of conspecifics.

We measured fitness related traits of singly mated-females housed at same-sex densities of 1, 2, or 10 and found clear reductions in lifetime reproductive output at low densities. Additionally, these females concentrated reproductive effort within early adulthood, albeit without improving estimates of fitness. When housed at variable densities, females altered their reproductive output in response to immediate densities, but were unable to improve it unless remating was possible, indicating an interaction between mating and density.

Overall, our findings suggest that reproductive plasticity in response to the presence of conspecifics mediates positive effects of density on fitness in female *Drosophila melanogaster*. Understanding the physiological and ecological bases of such plasticity may help explain the evolution of social tendency in the fly.

## 1. Introduction

Positive relationships between individual fitness and the abundance of individuals of any given species are referred to as Allee effects (Stephens et al., 1999). Such effects are in contrast to commonly observed negative density-dependent effects on population growth that result from resource competition or disease spread (Allee et al., 1949). A variety of mechanisms can underlie Allee effects, including demographic stochasticity in sex ratios, genetic mechanisms such as inbreeding depression, and inter-individual interactions (Courchamp et al., 1999; Stephens and Sutherland, 1999; Stephens et al., 1999).

Inter-individual interactions underlying Allee effects include examples of active co-operation, i.e. specialized interactions that yield collective fitness benefits, as well as passive phenomena, such as physiological facilitation or environmental modification (Allee et al., 1949; Courchamp et al., 1999). Such mechanisms are of particular interest to behaviour biologists, as they can help understand the evolution of co-operative interactions (Sachs et al., 2004) and explain patterns of sociality in a species (Stephens and Sutherland, 1999).

A growing body of work has demonstrated the existence of several inter-individual interactions in *Drosophila melanogaster*, including social aggregation (Navarro and del Solar, 1975; Simon et al., 2012), collective responses to food and threats (Tinette et al., 2004; Shultzaberger et al., 2019; Ramdya et al., 2015; Ferreira and Moita, 2020), non-random social networks (Schneider et al., 2012) and social learning (Sarin and Dukas, 2009; Mery et al., 2009; Kacsoh et al., 2015). The existence of such interactions strongly suggests that being near conspecifics may yield benefits, and consequently give rise to Allee effects.

Density-dependent population growth has been extensively studied in the fly (Pearl and Parker, 1922; discussed in Allee et al., 1949), but evidence in support of Allee effects is somewhat limited and variable. Pearl et al. (1927) demonstrated that lifespan increases with number of flies up to an optimal density between 35-55 individuals per bottle after which it declines. While the underlying mechanisms are not well understood, inter-individual interactions are associated with lifespan extensions in some mutant strains (Ruan and Wu, 2008) and may be responsible, in part, for such increases. Interestingly, other studies have reported reductions in lifespan with increases in adult density (Iliadi et al., 2009; Leech et al., 2017) suggesting that Allee effects on adult mortality may be context-specific.

Allee effects of larval density on larval survival (Sang, 1956; Ashburner et al., 2005) are typically thought to result from reduced feeding efficiency (Gregg et al., 1990; Dombrowski et al., 2017). Other factors such as risk of parasitism (Rohlfs and Hoffmeister, 2004) and competitive interactions with fungi (Rohlfs et al. 2005; Trienens and Rohlfs, 2020) are thought to strengthen Allee effects under natural conditions. However, as field studies have largely failed to find evidence of positive relationships between larval density and larval survival (Hoffmeister and Rohlfs, 2001; Wertheim et al., 2002), these effects are also likely to be context specific. Nevertheless, Allee effects on larval survival are commonly thought to bias oviposition site choice toward sites visited by other conspecifics (del Solar and Palomino, 1966). Consistent with this idea, the pheromone underlying gregarious oviposition, cis-vaccenyl acetate (cVA) (Wertheim et al., 2006; Dumenil et al., 2016) has been found to promote oviposition at intermediate pheromone concentrations, i.e. when larval densities may be high enough boost survival without incurring competition (Verschut et al., 2023).

An important mechanism by which density modulates individual fitness is via alteration of female reproductive output. By and large, effects of density on fecundity of female *Drosophila* have been found to be negative (Pearl and Parker, 1922; Chiang and Hodson, 1950; Barker, 1973; Rodriguez, 1989), which is consistent with reductions in fecundity following exposure to conspecific cues deposited on the food substrate (Fowler et al., 2022). However, a few studies have found evidence for Allee effects (Pearl, 1932; Robertson and Sang, 1944; Rockwell and Grossfield, 1978) which suggests that positive effects may be masked by strong negative effects under some experimental ecologies (Fujita 1954; Watt, 1960). Nutrient limitation due to crowding is a major factor determining the negative effects of density. However, as providing high-quality food in excess is not sufficient to observe positive effects of density in the fly (Robertson and Sang, 1944; Mueller and Huynh, 1994), we conjecture that other factors such as interference by males may also be involved. Increased proximity with males under laboratory conditions has measurable negative effects on female fertility (Linder and Rice, 2005; Orteiza et al., 2005; Edward et al., 2011) and lifespan (Fowler and Partridge, 1989). Given that most studies on density effects in the fly have used mixed-sex groups, negative effects of male presence may contribute to typically negative relationship seen between female fertility and density.

Assuming that density effects are additive, we may expect to observe Allee effects only when negative effects of density are controlled. To this end, we housed singly-mated females at non-crowding densities of 1, 2 or 10 individuals per vial (unpubl. data) without males, and examined their lifetime reproductive output, progeny survival and lifespan. We found clear evidence for an Allee effect as females housed in groups had higher reproductive progeny output than those housed singly or in pairs, regardless of genetic background. Females were able to modulate progeny production in response to changes in housing density but such plasticity failed to rescue lifetime reproductive output unless remating occurred. We also found some evidence for positive effects of adult density on larval survival. Taken together, we find strong support for the idea that individual fitness of female flies is enhanced by the presence of conspecifics.

## 2. Materials and Methods

### 2.1. Fly stocks

Two fly stocks were used for this study: Canton S and ChronoControls Merged (CCM). Canton S is an inbred strain maintained in vials at variable but small population sizes. By contrast, CCM flies are maintained as a single, randomly mating cage population with a population size of ∼2000 individuals (described in Gogna et al., 2015).

### 2.2. Fly maintenance

Parental flies were maintained in population cages as adults at densities of ∼600 and ∼2000 for Canton S and CCM respectively. Canton S flies were maintained in small cages (20.5 x 15.5 x 13.5 cm^3^) while CCM flies were kept in larger cages (25 × 20 × 15 cm^3^) due to their larger population sizes. To obtain age-matched experimental flies, flies were allowed to lay eggs on standard cornmeal medium containing charcoal for up to 12 hours, after which eggs were transferred to vial (diameter: 2.2 cm, height: 9.5 cm) without charcoal at densities of 60-70 per vial. Female pupae were collected on day-9 post egg collection and housed in fresh vials at different densities (see Section 3.3). Male pupae from the same vials were collected and maintained at densities of 10 or 20 per vial. All vials were maintained at 25°C under equinox light conditions (LD12:12).

### 2.3. Housing densities

All female pupae were housed in one of three housing densities - 1, 2 and 10 flies per vial- referred to as Single (S), Paired (P) or Grouped (G) henceforth (Figure 1). We chose a density of ten in the Grouped condition for logistical ease while also avoiding feeding competition (unpubl. data). We also included a paired treatment to distinguish between density-dependent and density-independent effects of conspecific presence, which can, in principle, act independently.

**Figure 1.**
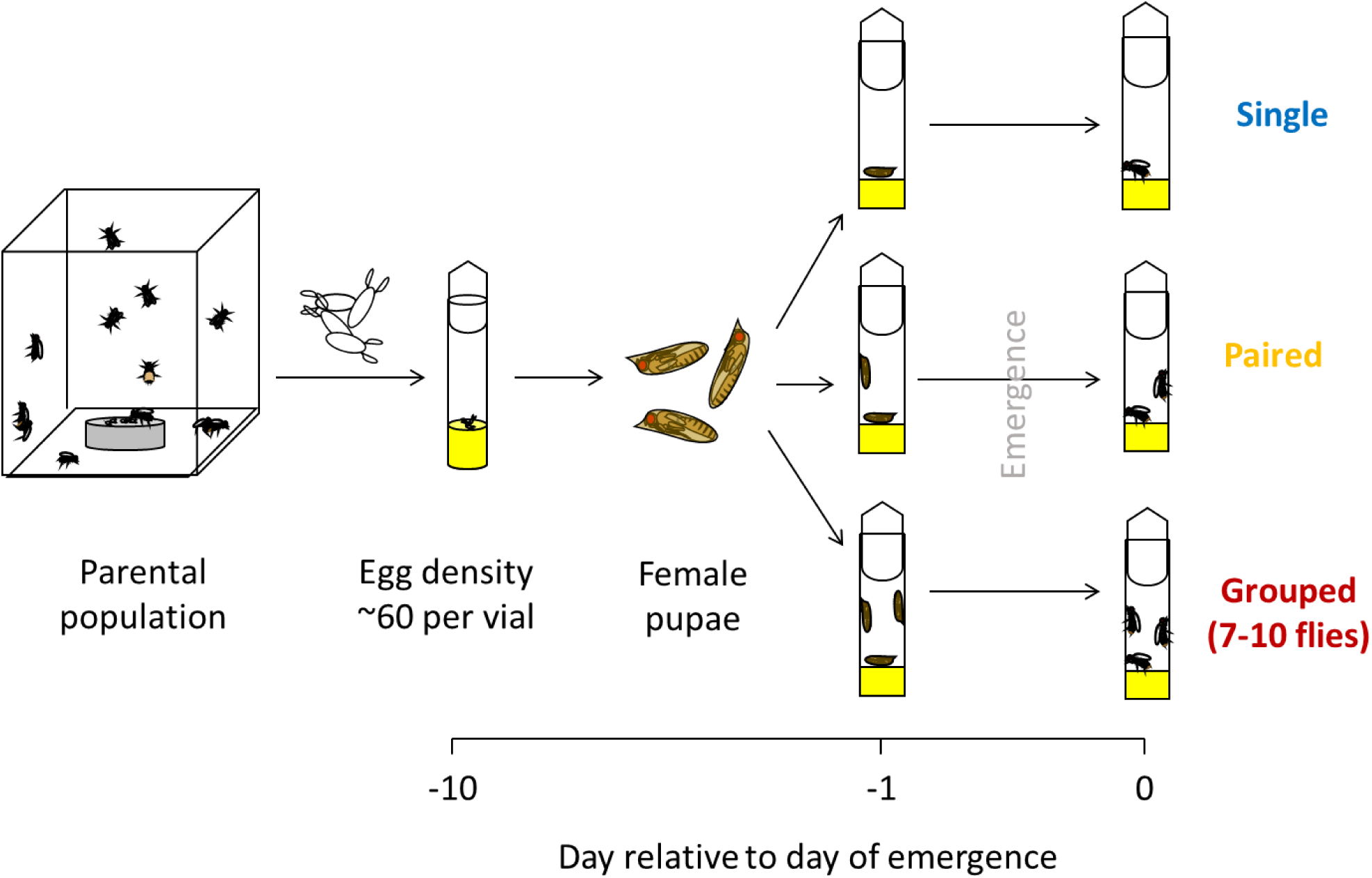
Protocol for generating experimental flies. Eggs laid over 12 hours were collected from parental cage populations of CCM or Canton S flies and cultured in vials containing standard cornmeal medium at densities of ∼60 eggs per vial. Female pupae were isolated from these vials on day 9 post egg collection and distributed across vials containing fresh cornmeal medium at different densities – 1 fly (Single), 2 flies (Paired) or 7-10 flies (Grouped).

As some pupae failed to eclose, we discarded all replicate vials with <70% of the expected number of flies. Across treatments, all vials contained exactly one mated female (see Section 3.4 for mating protocol). As all assays were conducted until the death of the mated female, densities varied over time due to mortality or losses during vial transfers. To ensure that mated flies experienced constant conspecific densities we replaced dead/lost companion flies with age-matched virgin females. As replacement flies were obtained from vials without mated females, fewer replacement flies were available towards the end of an assay. Consequently, grouped females had ≥7 companions over much of their reproductive phase (i.e. until cessation of progeny production), but reduced numbers closer to their death.

### 2.4. Mating protocol

On day-3 post emergence, single females were collected from each vial by aspiration and transferred to vials containing a thin layer of standard medium, referred to as mating vials hereafter. Vials were labelled to retain vial identity for each female and placed horizontally in a well-lit room maintained at 25°C for 30-60 min. At the end of this acclimatization period, a single male was aspirated into each mating vial. Mating behaviour was recorded manually by multiple observers, or using a camera for later observation. For each mating pair, the time of male introduction, start of copulation and termination of mating were recorded. These were used to calculate mating latency and duration.

### 2.5. Identity markings

For initial experiments, each mated female was transferred to a fresh vial containing her previous companions immediately after mating. Since the mated female could not be distinguished from virgins, the time of her death could not be measured. This was remedied in subsequent experiments by marking the thorax of a mated female post mating, using a black Staedler Lumocolor marker. Since flies cannot groom their thorax, these marks were retained throughout life. Since marking was performed under carbon dioxide anaesthesia, companion flies were also exposed to carbon dioxide prior to reunification with the mated female. This process was repeated in experiments where the females were remated, if the marks were found to be faint or otherwise difficult to detect.

### 2.6. Measuring components of fitness

To obtain the distribution of progeny across the female’s lifetime, all flies were transferred to fresh vials periodically, such that each vial contained eggs laid within a specific time-period. Transfers were performed daily for the first few days (7 or 10) post mating, and subsequently on every second day until the death of the mated fly (or all unmarked flies). As virgin companions laid unfertilized eggs which were indistinguishable from fertilized eggs laid by the mated female, reproductive output was measured as the number of progeny emerging from each vial instead of the total egg output. To facilitate larval feeding, particularly at low progeny densities, the food was dug with a scalpel before fly transfers. Vials were maintained at 25°C under LD12:12 conditions for at least 10 days after which adult progeny were counted.

The total number of progeny that emerged successfully represented the lifetime reproductive output for each mated female. Progeny found to be dead before emergence (i.e. as pupae or larvae) were also counted to estimate rates of pre-adult mortality, as the ratio of dead pre-adults to the total number of adult progeny (dead or live) found in the vial. It must be noted that as mortality during egg and early larval stages cannot be assessed accurately this protocol, these mortality values are likely to be underestimates. We measured lifespan of mated females by checking for marks on dead flies after each fly transfer and noting the ages at death for each marked individual.

To measure the combined effects of variation in different fitness-related traits, we estimated the intrinsic growth rate for each housing density. The intrinsic growth rate (λ) accounts for age-specific mortality and fertility rates to estimate the growth rate for a population upon reaching its stable age structure. We assumed that each treatment represents a population of females that ‘chooses’ its corresponding density as adults and is therefore subject to variation in fitness-related traits seen at that density. The λ, therefore, represents net fitness consequences associated with such a choice. λ values were measured for each treatment as dominant eigenvalue of Leslie matrices constructed using lifetime progeny counts obtained from the experiment. We employed a transformed Leslie matrix where survival probabilities were set to unity, and fertility values incorporated variation in pre-adult and adult mortality (Leslie, 1945). Such a transformed matrix has the same roots as the original form and yields the same estimate for λ (refer Supplementary methods for details).

### 2.7. Experimental design

Three sets of experiments were performed to test distinct sets of hypotheses, each following from previous results. Experiment 1 was performed with both Canton S and CCM flies, with qualitatively similar results in both cases. Hence, Experiments 2 and 3 were performed using only CCM flies. Lifespan and mating behaviour were measured only for the latter two experiments while pre-adult mortality rates were measured only for Experiment 1. Lifetime reproductive output was measured in all three experiments.

In Experiment 1, we tested if housing singly-mated females in isolation, pairs or groups of ten, influenced lifetime reproductive output and pre-adult mortality (Figure 2A). We expected to find higher values of fitness components for grouped flies if Allee effects were present, and a reverse relationship if only negative effects of density occurred.

**Figure 2.**
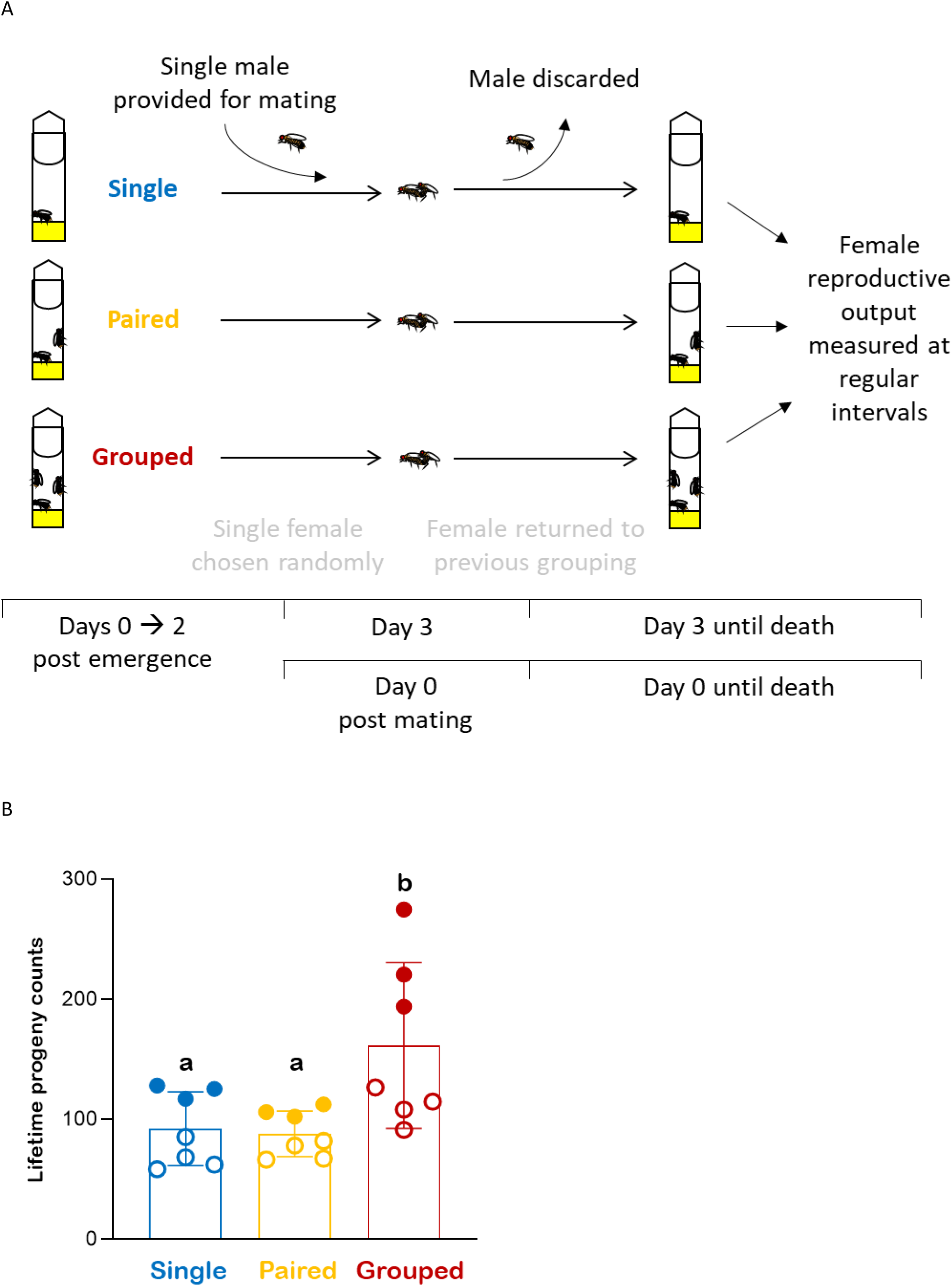
Housing density influences lifetime reproductive output. A) Schematic for Experiment 1- One female was chosen at random from each vial of the three treatments, and allowed to mate exactly once before being returned to a fresh vial with her previous companions, if any. Subsequently, all flies were transferred to fresh vials daily for the first seven days, after which they were transferred every two days until all females died. Used vials containing eggs laid since the previous transfer were kept under LD 12:12 conditions at 25°C until all adult progeny had emerged and were counted. Females were mated on the 3^rd^ day post emergence. All subsequent days were dated with the day of mating as a reference. Flies from Canton S and CCM genetic backgrounds were used for this experiment. B) Grouped flies have higher lifetime reproductive output, measured as the total number of live adult progeny produced by the focal female, compared to singly housed and paired flies. Scatter points represent mean values for N replicate experiments, each obtained by averaging data across *n* individuals within the replicate. Open and closed circles indicate replicates with Canton S and CCM flies respectively. Height of each bar reflects the mean across N replicate experiments while error bars indicate the corresponding standard deviation. Bars which share any letter are not significantly different from one another. Data analysed using a two-way ANOVA with Housing density and Strain as fixed factors. Pairwise comparisons performed using post-hoc Tukey’s HSD tests (N = 4 and 3 for Canton S and CCM respectively, *n* = 8 - 10 per treatment).

**Figure 3.**
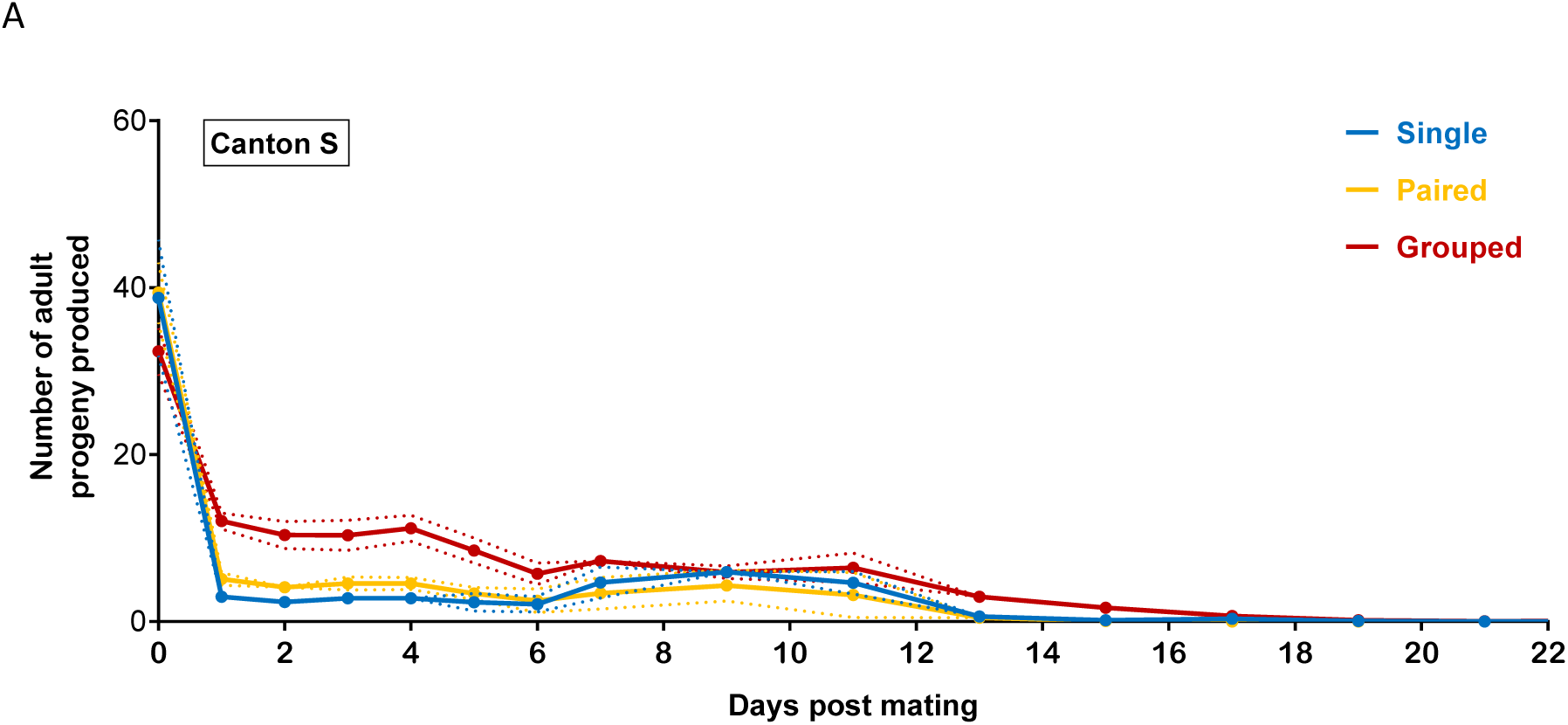
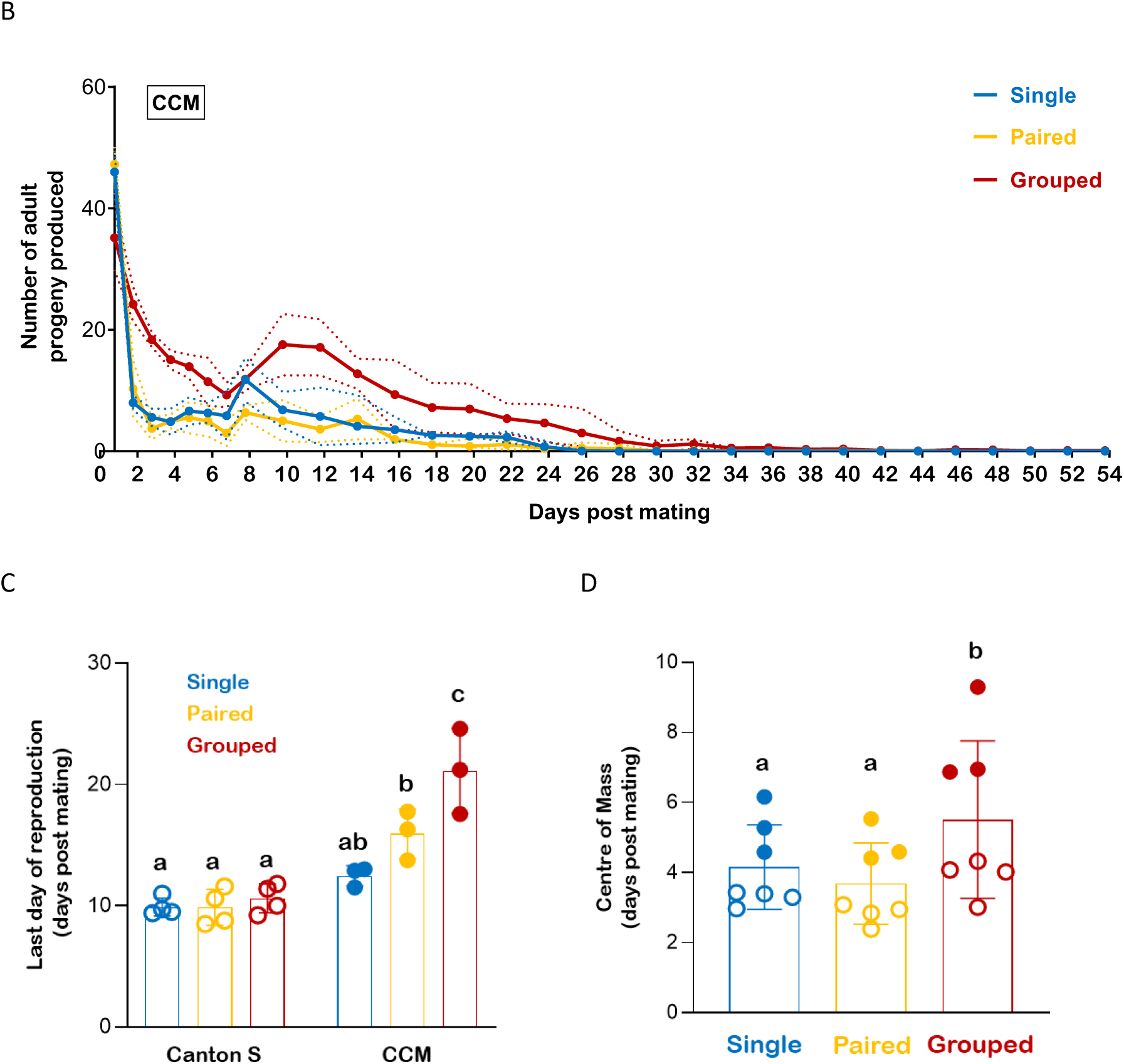
Distribution of progeny numbers across the lifetime is modified by housing density. Patterns of progeny output across time are different for grouped flies compared to singly housed and paired female in (A) Canton S and (B) CCM backgrounds. Note that the increase in progeny counts after day 7 is due to pooling of progeny output over two days. Data averaged across *n* individuals within each replicate experiment and subsequently across N replicate experiments. Dotted lines indicate standard deviation in progeny counts across N replicate experiments. (N = 4 and 3 for Canton S and CCM respectively, *n* varied between 8 - 10 per treatment). Grouped flies show delayed patterns of reproduction, as is evident from (C) the last day of reproduction, reflecting the period of successful reproductive activity for a female, and (D) centre of mass (CoM) of the distribution of progeny across time. Scatter points represent mean values for N replicate experiments, each obtained by averaging data across *n* individuals within the replicate. Open and closed circles indicate replicates with Canton S and CCM flies respectively. Height of each bar reflects the mean across N replicate experiments while error bars indicate the corresponding standard deviation. Bars which share any letter are not significantly different from one another. Data analysed using a two-way ANOVA with Housing density and Strain as fixed factors. Pairwise comparisons performed using post-hoc Tukey’s HSD tests (N = 4 and 3 for Canton S and CCM respectively, *n* = 8 - 10 per treatment).

Since freely moving flies can modulate the number of flies around them, we hypothesized that female reproductive output depends only on immediate conspecific densities. In Experiment 2, we examined the role of density-dependent reproductive plasticity in mediating differences in lifetime reproductive output across treatments by varying immediate densities for singly-mated females over adulthood. Flies were housed, as above, for three days post mating, before being transferred to a different density for the rest of their lives (Figure 6A). Previously grouped flies were either maintained in groups (GG), or transferred into pairs (GP) or single housing (GS). Previously paired or single housed flies were maintained in their previous group size (PP or SS) or transferred to groups (PG or SG respectively). Owing to logistical constraints, we did not include treatments where single flies were transferred into pairs and vice versa (SP and PS).

**Figure 4.**
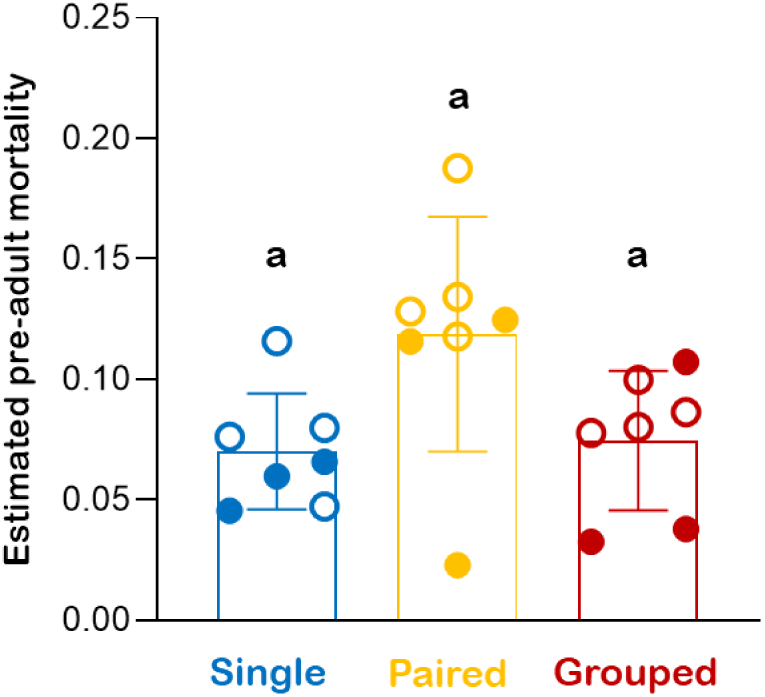
Adult density influences progeny mortality during the pre-adult stages. Progeny mortality during the pre-adult stages, estimated as the fraction of live adult progeny produced by the focal female, varies significantly across housing densities (See Section 4.3 for details). Mortality rates across days and individuals were sorted according to their associated egg density and mean mortality for different egg densities was calculated. Scatter points represent mean values for N replicate experiments obtained after averaging across these egg densities within each replicate. Open and closed circles indicate replicates with Canton S and CCM flies respectively. Height of each bar reflects the mean across N replicate experiments while error bars indicate the corresponding standard deviation. Bars which share any letter are not significantly different from one another. Data analysed using a repeated measures ANOVA with Housing density and Strain as fixed factors and Egg density as the repeated measures factor. Pairwise comparisons performed using post-hoc Tukey’s HSD tests (N = 4 and 3 for Canton S and CCM respectively, *n* = 8 – 10 per treatment).

**Figure 5.**
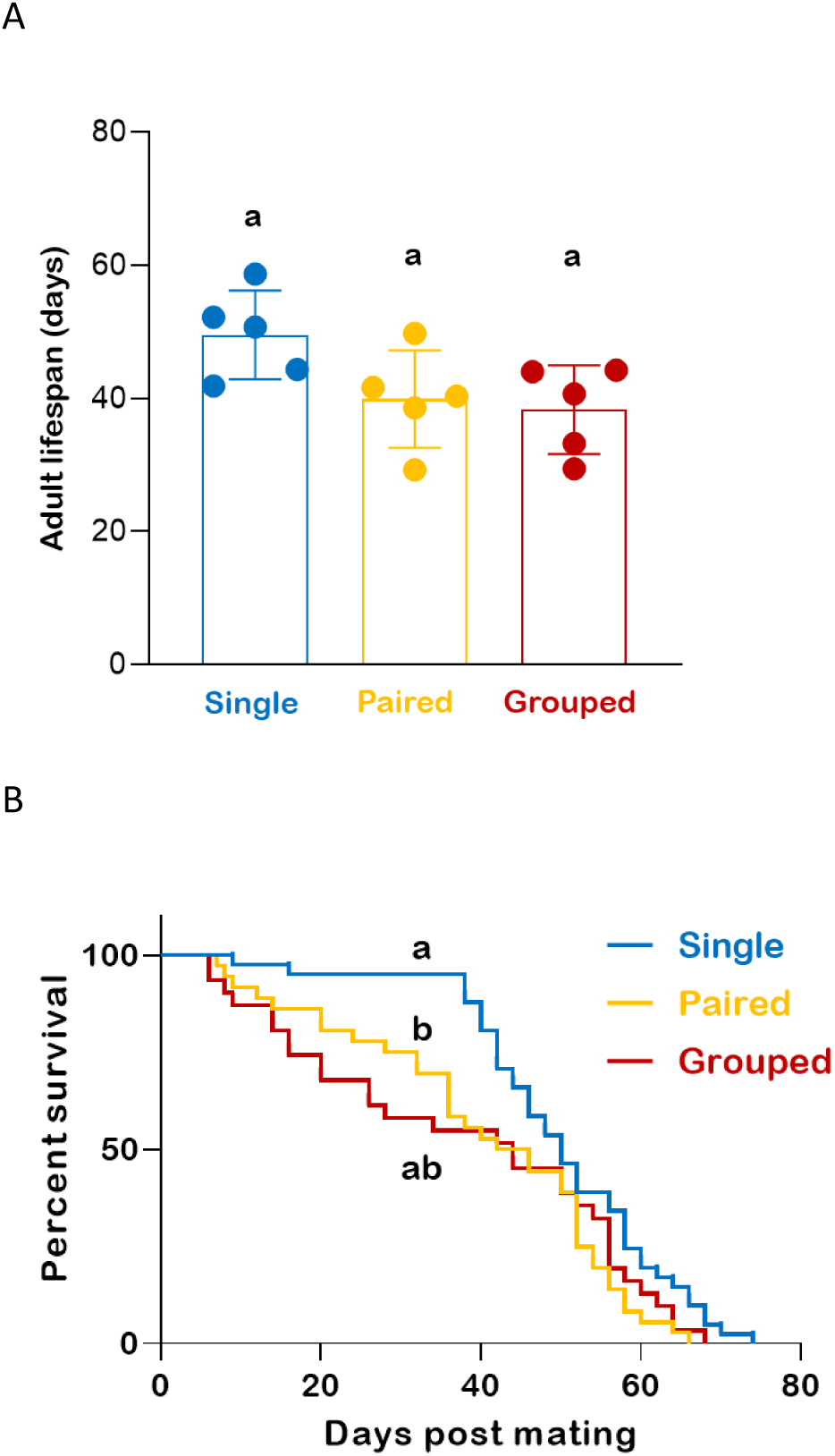
Exposure to conspecifics reduces survival of the mated female. A) Mean adult lifespan of the mated female does not vary significantly across housing densities. Scatter points represent mean values for N replicate experiments, each obtained by averaging data across *n* individuals within the replicate. Height of each bar reflects the mean across N replicate experiments while error bars indicate the corresponding standard deviation. Bars which share any letter are not significantly different from one another. Data analysed using a mixed-model ANOVA with Housing density as a fixed factor and Replicate as a random factor. Pairwise comparisons performed using post-hoc Tukey’s HSD tests (N = 5 and *n* = 4 - 10 per treatment). B) Singly housed flies experience higher survival during early adulthood, as seen from survival curves for mated females obtained by pooling individuals across 5 replicate experiments (*n* = 31 - 41). Log-rank tests were used for omnibus testing and for pairwise comparisons. Curves which share any letter are not significantly different from one another.

**Figure 6.**
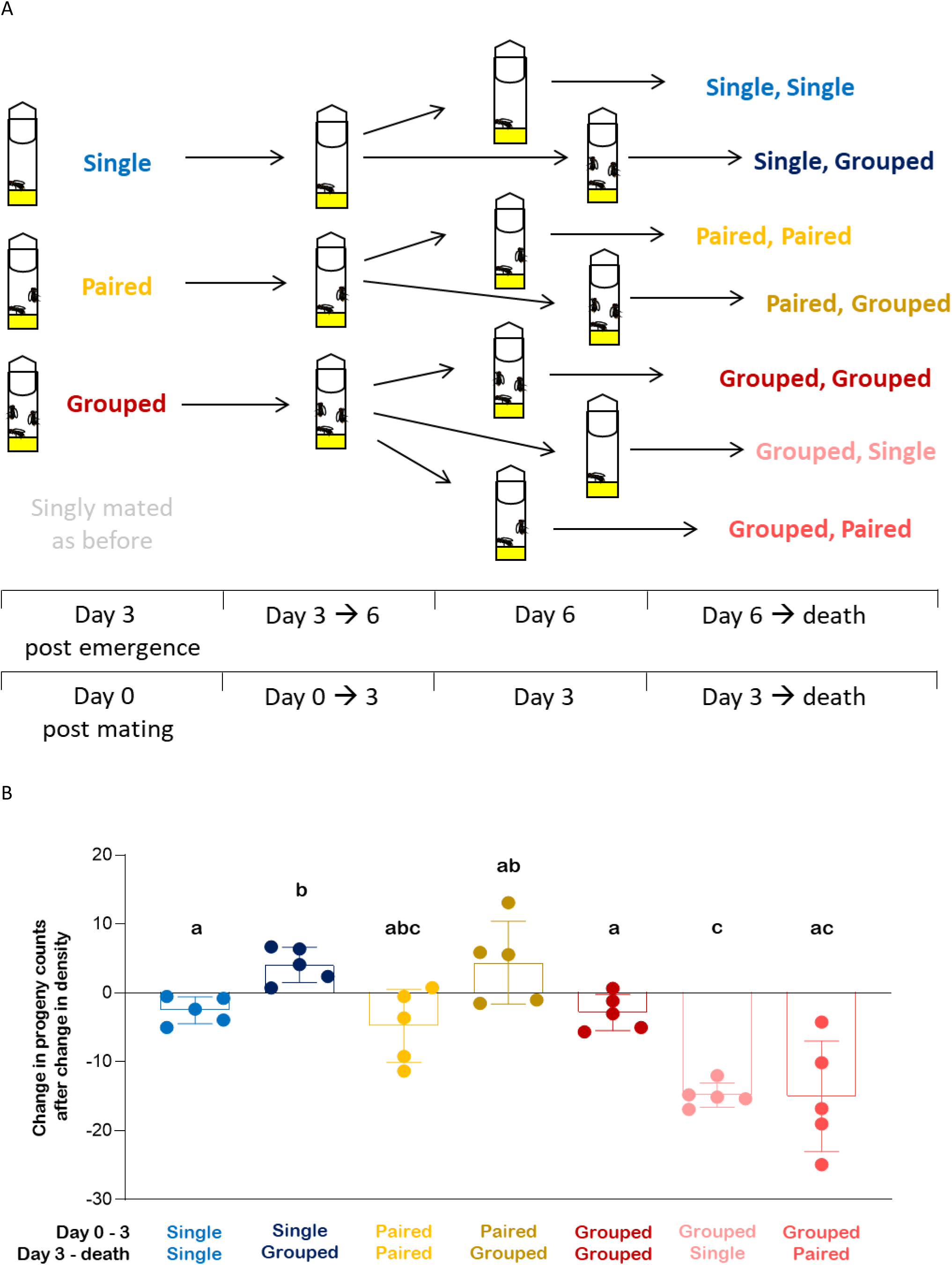
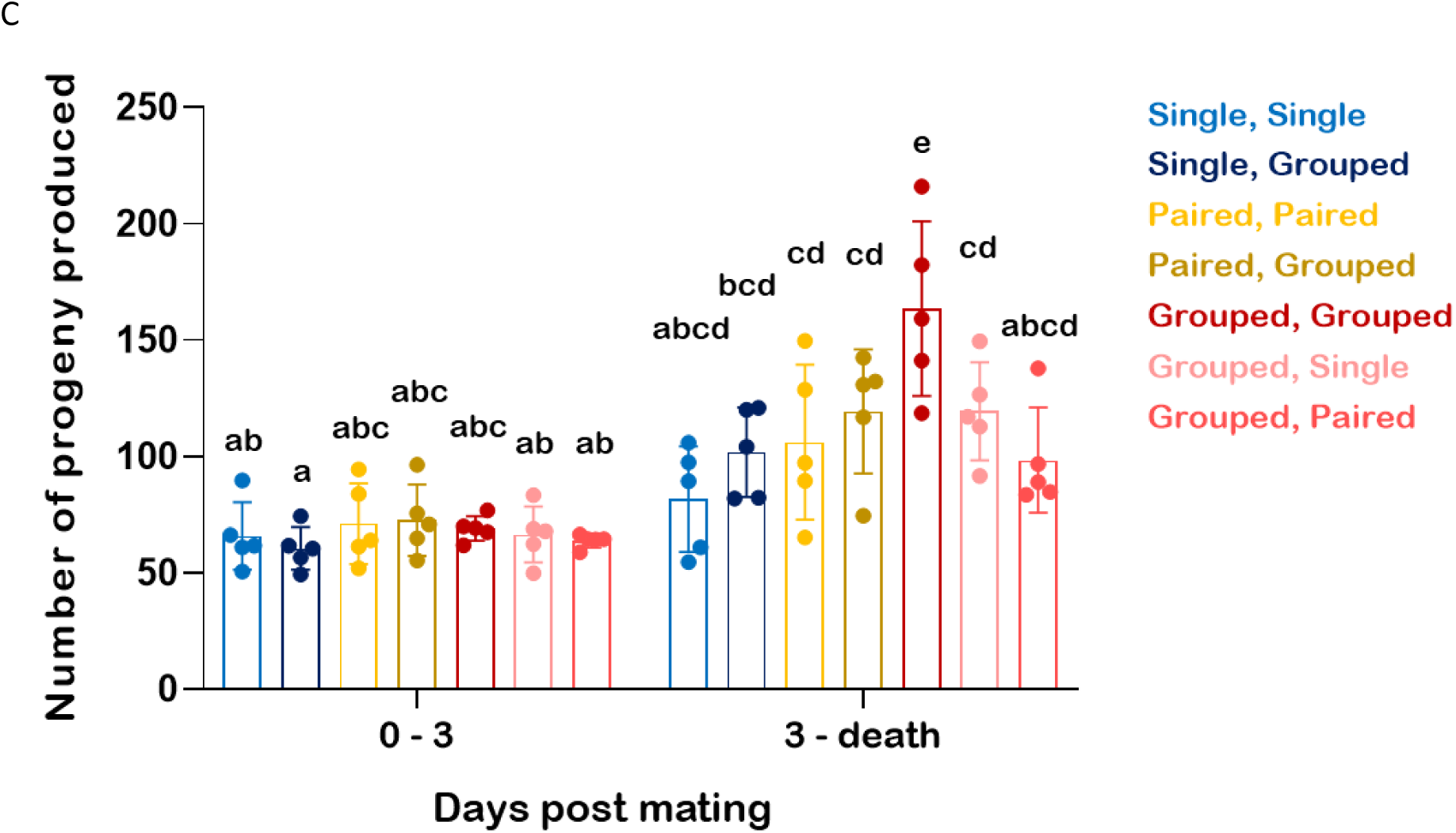
Immediate and previously experienced conspecific densities influence progeny output of singly mated females. A) Schematic for Experiment 2- Single females from each vial were mated and housed as in Experiment 1 until day 3 post mating. On day 3, subsets of flies from each density were transferred to vials with a different density, as indicated. All flies were transferred to fresh vials daily for the next seven days, after which they were transferred every two days until all females died. Used vials containing eggs that were laid since the previous transfer were kept under LD 12:12 conditions at 25°C until all adult progeny had emerged and were counted. Females were mated on the 3^rd^ day post emergence and all subsequent days were dated with this day as a reference. Only flies from the CCM genetic background were used for this experiment. B) Females increase or decrease their immediate progeny output in response to increases or decreases in housing densities. Change in progeny production in response to changes in density was measured as the difference between progeny output for days 3 and 4 post mating. C) Plastic changes in progeny output fail to modify lifetime reproductive output if females have previously experienced low densities. Scatter points represent mean values for N replicate experiments, each obtained by averaging data across *n* individuals within the replicate. Height of each bar reflects the mean across N replicate experiments while error bars indicate the corresponding standard deviation. Bars which share any letter are not significantly different from one another. For B), data were analysed using a Welch test with Housing condition as the factor of interest, and Games-Howell tests for pairwise comparisons (N = 5, *n* = 6 - 10 per treatment). For C), data were analysed using a repeated measures ANOVA with Housing condition as the fixed factor and Phase of the experiment as the repeated measure, and post-hoc Tukey’s HSD tests for pairwise comparisons (N = 5, *n* = 4 - 10 per treatment).

In Experiment 3, we examined if reproductive plasticity was influenced by the age of exposure to conspecifics and remating. To assess the effect of age, we varied the age at which a female experienced single housing before being housed in a group for the rest of her life. For this we housed singly-mated females in one of three treatments– continuous group housing (GGG), single housing until day-3 post mating (SGG) or single housing from day-3 to day-6 post mating (GSG) (Figure 7A). Note that in each case, the mated female was housed in groups after day-6 post mating. To test the effect of remating, we provided the mated female with a single remating on day-8.

**Figure 7.**
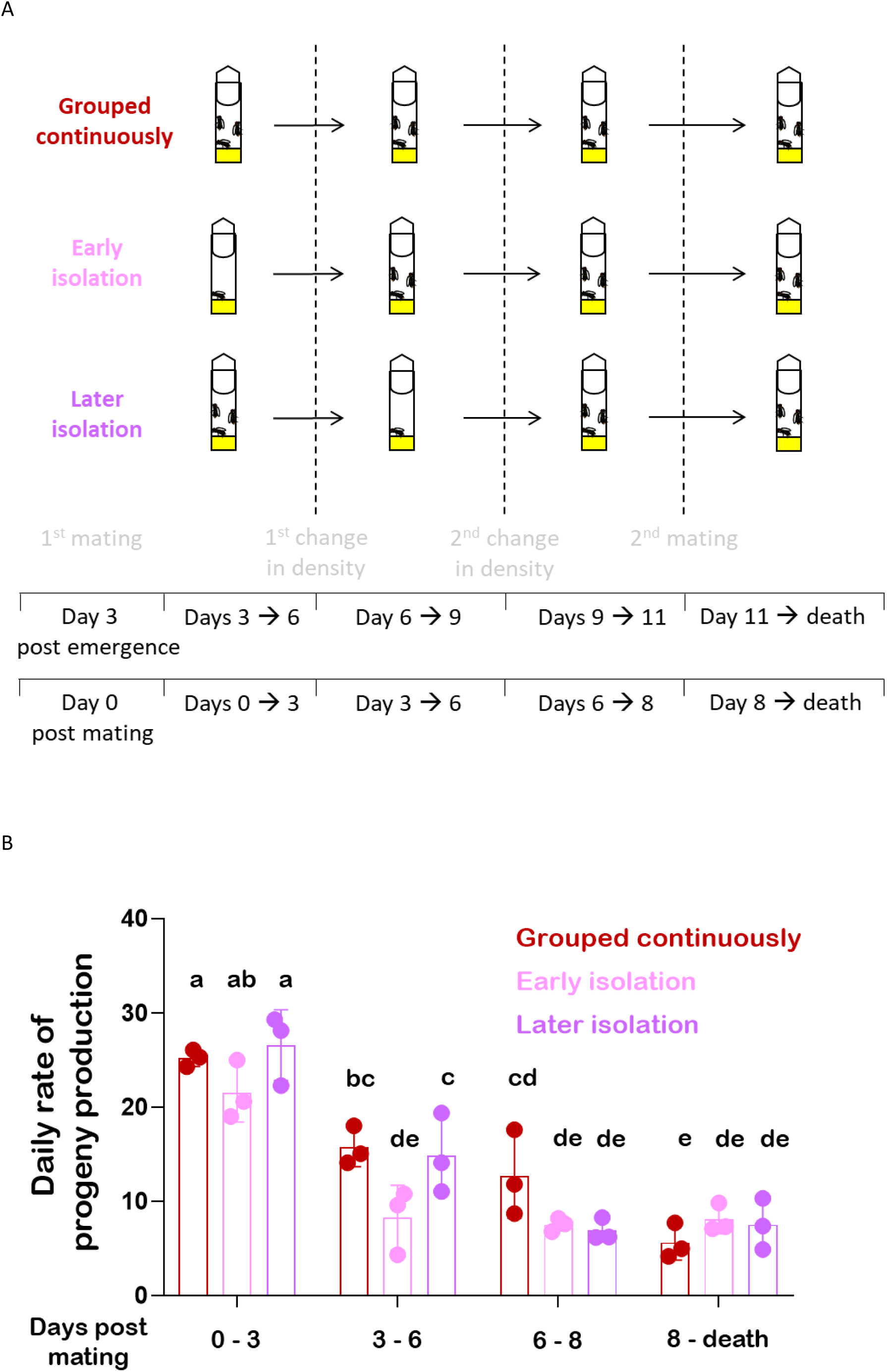
Immediate and previously experienced conspecific densities influence progeny output of singly mated females. A) Schematic for Experiment 3- Females were housed either singly or in groups and treated as in Experiment 1 until day 3 post mating. On day 3, all singly housed females (Early isolation) were transferred to vials with groups of virgin females and housed thus for the rest of the assay. Half of the grouped females were housed singly between days 3 and 6 (Later isolation) before rejoining their old groups, while the other half remained in groups throughout their lives (Grouped continuously). Note that all mated females remained in groups after day 6. On day 8, all mated females were allowed to remate once. All flies were transferred to fresh vials daily for the next seven days, and every two days thereafter, until all females died. Used vials containing eggs that were laid since the previous transfer were kept under LD 12:12 conditions at 25°C until all adult progeny had emerged and were counted. Females were mated on the 3^rd^ day post emergence and all subsequent days were dated with this day as a reference. Only flies from the CCM genetic background were used for this experiment. B) Daily rates of progeny production are lower for females that previously experienced low densities compared to females that remained in groups continuously. These differences are lost after remating. Scatter points represent mean values for N replicate experiments, each obtained by averaging data across *n* individuals within the replicate. Height of each bar reflects the mean across N replicate experiments while error bars indicate the corresponding standard deviation. Bars which share any letter are not significantly different from one another. Data analysed using a repeated measures ANOVA with Housing condition as a fixed factor and Phase of the experiment as the repeated measure. Post-hoc Tukey’s HSD tests were used for pairwise comparisons (N = 3, *n* = 5 - 9 per treatment).

### 2.8. Statistical analyses

We performed each experiment in replicate blocks for logistical ease, with each replicate containing ten singly-mated females per treatment. Since flies were often lost over the course of the experiment, there was variation in the number of usable individuals across treatments and replicates. Hence, we used replicate as our level of replication and averaged data across individuals within a replicate per treatment. Most analyses employed mixed-model ANOVAs with Replicate as a random factor. We verified ANOVA assumptions using Levene’s tests and Q-Q plots. In cases of violations, data were appropriately transformed before analyses. Non-parametric tests were used when data transformation failed to correct for the violation of assumptions.

Intrinsic growth rates (λ) were estimated as eigenvalues of Leslie matrices constructed using mean lifetime progeny counts (pooled across replicates) (Leslie, 1945) for different density treatments. Hypotheses about these growth rates were tested using randomization methods (refer Supplementary methods for details). To obtain a randomized dataset, individual-level data were randomly shuffled across treatments before averaging counts for each treatment. These randomized count means yielded randomized λ values for each treatment. For hypothesis testing, observed test statistics were compared to sets of 1000 randomized values and fractions of randomized values greater than or equal to observed values were calculated as p-values. Benjamini-Hochberg corrections with a false discovery rate of 0.05 were applied when sets of hypotheses were tested.

Parametric analyses were performed using Statistica v.7 while randomization analyses were performed using custom scripts in MATLAB 2022b. Welch’s tests and Benjamini-Hochberg corrections were performed using resources provided by McDonald (2014). Log rank tests for comparing survival curves were performed using GraphPad Prism 8. All figures were prepared using GraphPad Prism 8 and MS PowerPoint. All schematics were prepared using MS PowerPoint.

## 3. Results

### 3.1. Females housed in groups have higher lifetime reproductive output than females housed singly or in pairs

Comparison of total progeny counts across treatments in Experiment 1 revealed that number of conspecifics modifies lifetime reproductive output in females from both Canton S (Main effect of Conspecific number: *F_2,6_* =20.399, *p*=0.002) and CCM (Main effect of Conspecific number on log transformed values: *F_2,4_* =82.029, *p*=0.0006) backgrounds. Post-hoc tests showed that this effect was due to significantly higher lifetime progeny counts for females housed in groups compared to those alone or in pairs. Similar results were obtained when genotype was included as a fixed factor (Main effect of Conspecific number on log transformed values: *F_2,15_* =44.46, *p* < 10^-6^) (Figure 2B).

These data suggest that lifetime reproduction is enhanced by the presence of multiple conspecifics, but not a single conspecific (i.e. paired housing).

### 3.2. Distribution of progeny across time varies with conspecific density

In addition to differences in progeny number, we observed considerable variation in the distribution of progeny across a female’s lifetime (Figure 3A-B). Given that timing of reproduction can influence fitness (Brommer, 2000), we characterized these differences further. Qualitative assessment of distributions across individuals indicated that grouped flies had continuous patterns of progeny production across time, while single and paired flies produced the bulk of their progeny on the first day and the rest in brief bouts (Supplementary figure 1A-C).

To describe such variation quantitatively, we measured the widths and centres of mass (CoM) for progeny distributions across treatments. The width represents the duration of progeny production, or the reproductive period, and is determined by the last day on which viable progeny were produced. The CoM reflects the mean age of progeny production, calculated as the weighted sum of days lived by a female with daily progeny fractions as weights. Small values of CoM suggest that progeny output is biased towards early ages while larger values suggest that it is delayed.

We found that the last day of reproduction did not differ significantly across densities in Canton S flies (Main effect of Conspecific number: *F_2,6_* =0.534, *p*=0.61) but was significantly delayed for CCM flies housed in groups (Main effect of Conspecific number: *F_2,4_* =7.397, *p*=0.045). This was reflected in a significant Genotype x Conspecific number interaction when data were pooled across genotypes (Effect of Genotype x Conspecific number: *F_2,15_* =8.683, *p* =0.0031) (Figure 3C). Significant effects of conspecific densities on CoM were detected for both Canton S (Main effect of Conspecific number on log transformed values: *F_2,6_* =30.05, *p*=0.0007) and CCM (Main effect of Conspecific number on square root transformed values: *F_2,4_* =9.088, *p*=0.032) flies. Similar results were obtained when genotype was included as a fixed factor (Main effect of Conspecific number on square root transformed values: *F_2,15_* =15.692, *p*=0.0002). Post hoc tests showed that the CoM was significantly delayed in grouped flies compared to both single housed and paired flies in both genotypes (Figure 3D).

Taken together, females appear to vary their reproductive tactic (Emlen, 2008) in response to conspecific presence– at low densities, reproductive activity is concentrated in the period immediately after mating, while at higher densities, progeny are produced at a low, consistent rate.

### 3.3. Adult density but not larval density influences larval survival

Given that progeny output varied across treatments, adult density could influence pre-adult survival indirectly, via its effects on progeny numbers. To separate these indirect effects from any direct effects of adult density on pre-adult survival, we sorted pre-adult mortality data by the estimated pre-adult density in each vial, and obtained mean mortality values for different pre-adult densities within each replicate experiment.

Analysis of these data with genotype and adult density as fixed factors, and estimated pre-adult density as a within-replicates factor revealed a significant effect of adult density (Main effect of Conspecific number: *F_2,15_* = 4.345, *p*=0.032) (Figure 4) but no significant effect of pre-adult density on pre-adult survival (Main effect of pre-adult density: *F_3,45_* = 3.242, *p*=0.031; After Greenhouse-Geiser correction: *F_1.4,21.3_* = 3.242, *p*=0.073) (Supplementary figure 2). Although pairwise comparisons among the treatments did not yield significant differences, visual examination suggests that progeny of paired females experienced higher mortality than either singly housed or grouped females.

Taken together, these data suggest that the number of adult females present on the oviposition site can influence pre-adult survival.

### 3.4. Variation in total reproductive output is largely responsible for variation in intrinsic growth rates across conspecific densities

The above results suggest that different fitness-related traits are maximized at different adult densities. While grouped flies have the highest total reproductive output (Figure 2B), single and paired flies reproduce earlier (Figure 3D) which may provide them with head start advantages. Additionally, pre-adult survival varies across densities (Figure 4). Thus, to identify the density at which overall fitness is maximized, we used intrinsic growth rates as a composite measure of variation in age-specific fertility and mortality.

For both genotypes, the observed variation in λ across adult densities (Table 1) was significantly greater than expected by chance alone (Canton S: *p*=0.006; CCM: *p*= 0) suggesting that variation in fitness components across densities is associated with significant changes in the overall growth rate. Pairwise comparisons revealed that grouped flies had significantly larger λ than singly housed (Canton S: *p*=0; CCM: *p*= 0) and paired flies (Canton S: *p*=0.046; note that this difference was not significant after BH correction: *p*=0.069; CCM: *p*= 0). However, differences between single and paired flies were not significant for either genotype (Canton S: *p*=0.132; CCM: *p*=0.414).

**Table 1.** Intrinsic growth rates (λ) vary across housing densities. Grouped flies have higher intrinsic growth rates, estimated as the dominant eigenvalue of a Leslie matrix constructed using the corresponding progeny count data across time, compared to singly housed and paired flies. Separate estimates were obtained for Canton S and CCM flies and compared within each genetic background using randomization tests. Rows which share any letter are not significantly different from one another.

| Strain | Treatment | Estimate of $\lambda$ | Pairwise difference |
| --- | --- | --- | --- |
| Canton S | Single | 1.275 | a |
|  | Paired | 1.286 | ab |
|  | Grouped | 1.300 | b |
| CCM | Single | 1.307 | a |
|  | Paired | 1.306 | a |
|  | Grouped | 1.334 | b |

To assess contribution of each fitness component to λ, we systematically randomized mortality and fertility data across treatments and compared the resultant λ values to the observed ones (refer Supplementary methods for details). If the observed λ differed significantly from that obtained after a given component was randomized, then that component was considered necessary to produce the observed λ. If the observed λ was significantly different from that obtained when a component was selectively not randomized, then the component was considered insufficient.

Total reproductive output, but not the distribution of progeny across time, was found to be necessary to produce the observed λ values for singly housed flies from both genotypes (Table 2). Thus, contrary to expectation, advancement of reproduction failed to improve growth rates for these flies. Similar patterns were seen for paired flies from both genotypes, although necessity of total reproductive output in the Canton S strain had limited statistical support (*p*=0.063).

**Table 2.** Contribution of different components of fitness to intrinsic growth rate (λ) varies across housing densities. Variation in lifetime reproductive output largely determines variation in intrinsic growth rates (λ) across housing densities. Table 2 compares λ values estimated from the observed data to λ values expected under different null hypotheses of sufficiency and necessity for individual components of fitness – lifetime reproductive output which represents the total number of progeny produced regardless of their survival to adulthood, distribution of reproductive output across age, and maternal age-specific mortality rates for progeny. Values in brackets represent *p*-values under a given null hypothesis, obtained by calculating the fraction of randomized values more extreme than the observed value and applying a Benjamini-Hochberg correction for multiple comparisons. Cases where the null hypothesis is rejected are highlighted in red.

| Treatment | Component of fitness | Canton S |  |  | CCM |  |  |
| --- | --- | --- | --- | --- | --- | --- | --- |
| | | Observed $\lambda$ | Expected $\lambda$ under $H_0$ of | | Observed $\lambda$ | Expected $\lambda$ under $H_0$ of | |
|  |  |  | Sufficiency | Lack of Necessity |  | Sufficiency | Lack of Necessity |
| Single | Lifetime reproductive output | 1.275 | 1.277<br>(0.622) | 1.304<br>(0) | 1.307 | 1.307<br>(0.993) | 1.338<br>(0) |
|  | Distribution of progeny across time |  | 1.306<br>(0) | 1.278<br>(0.459) |  | 1.339<br>(0) | 1.307<br>(0.998) |
|  | Pre-adult mortality |  | 1.288<br>(0) | 1.275<br>(0.632) |  | 1.317<br>(0.015) | 1.308<br>(0.683) |
| Paired | Lifetime reproductive output | 1.286 | 1.287<br>(0.794) | 1.303<br>(0.063) | 1.306 | 1.297<br>(0.151) | 1.345<br>(0) |
|  | Distribution of progeny across time |  | 1.305<br>(0.028) | 1.287<br>(0.738) |  | 1.347<br>(0) | 1.296<br>(0.084) |
|  | Pre-adult mortality |  | 1.287<br>(0.852) | 1.287<br>(0.507) |  | 1.316<br>(0.023) | 1.307<br>(0.161) |
| Grouped | Lifetime reproductive output | 1.300 | 1.324<br>(0) | 1.288<br>(0.13) | 1.334 | 1.367<br>(0) | 1.309<br>(0.004) |
|  | Distribution of progeny across time |  | 1.287<br>(0.103) | 1.326<br>(0) |  | 1.307<br>(0) | 1.369<br>(0) |
|  | Pre-adult mortality |  | 1.289<br>(0.016) | 1.3<br>(0.622) |  | 1.319<br>(0) | 1.332<br>(0.036) |

In case of grouped flies, high reproductive output alone yielded significantly greater growth rates than observed (i.e. was insufficient to produce the observed λ). Smaller observed values were attributable to the effects of delayed reproduction, suggesting that both reproductive output and the distribution of progeny production are necessary to for observed λ. Such a pattern was evident in both genotypes but had statistical support for only CCM flies (Effect of reproductive output: *p_CS_* = *p_CCM_* = 0; Effect of progeny distribution: *p_CS_* = 0.1, *p_CCM_* = 0).

Interestingly, pre-adult mortality rates failed to contribute significantly to growth rates in all sets of flies except grouped CCM flies.

Taken together, these data suggest that low growth rates in singly housed and paired flies are primarily due to the effects of low reproductive output which cannot be overcome by advancement of reproductive activity. By contrast, delayed reproduction serves to diminish growth rates of grouped flies which is compensated for by higher reproductive output.

### 3.5. Female lifespan is influenced by the presence of conspecifics

As mated flies were distinguishable from their companions in Experiment 2, we were able to assess time of death and hence measure lifespan for these females across treatments.

Although we failed to find statistically significant associations between lifespan of the mated female and conspecific density (Main effect of social environment: *F_2,8_* =2.693, *p*=0.128), comparison of survival curves indicated significant variation across densities (*Χ^2^_df=2_* =6.944, *p*= 0.031) (Figure 5A-B). Singly housed flies lived longer than both paired and grouped flies, although statistical significance was detected only with the former (*Χ^2^_df=1_* =6.433, *p*= 0.011). Such trends are consistent with previous reports (Iliadi et al. 2009; Leech et al., 2017). Notably, survival curves and mean lifespan were both largely comparable for paired and grouped flies, suggesting that conspecific presence may have density-independent effects on lifespan of the mated female.

### 3.6. Lifetime reproductive output is sensitive to immediate conspecific density as well as previously experienced densities

As differences in reproductive output across densities are evident immediately after mating, we considered that females might evaluate conspecific numbers within their immediate environment and modulate progeny output accordingly. Under such a scenario, changes in density are expected to elicit immediate changes in the rate of progeny production. Moreover, if differences in lifetime reproductive output result from oviposition choices of the female, such plasticity may allow females to reconsider their choices and recover their reproductive output. We tested this hypothesis in Experiment 2 by comparing progeny output of females that underwent a change in the number of their companions and those that remained at constant housing densities.

When immediate changes in progeny counts were measured (difference in progeny counts between day-2 and day-1 after changing densities), single flies were found to exhibit significant increases upon grouping while grouped flies showed significant reductions upon isolation (Main effect of Treatment: *F_6,12.2_*= 32.27, *p*=8.8 x 10^-7^; Games-Howell tests: *p*<0.05) (Figure 6B). Similar trends were observed when flies were transferred to and from pairs, although they failed to reach statistical significance. These results suggest that female flies can assess their immediate conspecific density and modulate their progeny output accordingly.

However, when we compared the total progeny output after changes in density, we found that plastic responses to the altered density did not necessarily influence lifetime reproduction (Figure 6C). Flies that experienced single or paired housing failed to show higher progeny output after conspecific numbers were increased. By contrast, previously grouped flies showed significant reductions in progeny output after conspecific numbers were reduced (Effect of Treatment x Phase of the experiment: *F_6,28_*= 5.394, *p*=0.0008; Tukey’s HSD tests: *p*<0.05).

Thus, while females are capable of modulating progeny output in response to immediate density, such modulation fails to rescue negative effects of previously experienced low densities. In contrast to such long-lived effects of low-density exposure, effects of exposure to higher densities failed to persist after densities were reduced.

### 3.7. Effects of low conspecific exposure are lost after remating

Since behavioural maturation occurs during early adulthood, we considered that the inability of singly housed flies to recover progeny production after grouping may be due to the absence of conspecifics during a developmentally critical period for reproductive behaviour. Consequently, long-lived effects of single housing may be expected to be absent at later ages. We tested this hypothesis in Experiment 3 and failed to find statistically significant differences in the rates of progeny production among flies that experienced single housing at different ages and those housed in groups (Effect of Treatment x Phase of the experiment: *F_6,18_*= 4.86, *p*=0.004; Tukey’s HSD tests: *p*>0.05) (Figure 7B, Day-6 - 8). It must be noted, however, that grouped flies tended to have higher rates of progeny production than flies that had experienced isolation at any age. While this warrants further study, we speculate that developmental effects do not underlie the inability of single housed flies to recover progeny output after grouping.

An alternative explanation for the inability of singly-housed flies to recover progeny output after grouping may be that these flies carry few sperm by the time they experience permissible conspecific densities. If this were true, then we would expect such females to recover progeny production if allowed to remate and replenish sperm reserves. Consistent with this idea, daily rates of progeny production were comparable for females with and without prior single housing after they were allowed to remate (Effect of Social environment x Phase of the experiment: *F_6,18_*= 4.86, *p*=0.004; Tukey’s HSD tests: *p*<0.05) (Figure 7B, Day-8 - death). This is markedly unlike Experiment 2, where singly housed females failed to match progeny output of continuously grouped females after being grouped (Figure 6B, Day-3 – death).

Overall, these results suggest that variation in progeny output may result from density-dependent variation in a female’s sperm reserves rather than developmental effects.

### 3.8. Mating behaviour is unaffected by presence of conspecifics

Variation in sperm reserves across housing densities could result from differences in sperm storage in the female reproductive tract after mating or differential male investment during mating itself. To test the latter hypothesis, we examined behaviour of mating pairs from Experiments 2 and 3. Males are known to increase copulation duration to transfer greater amounts of accessory gland proteins (but not sperm itself) which can influence sperm storage (Gilchrist and Partridge, 2000; Tram and Wolfner, 1999). Hence, we expected to find differences in copulation duration if variation in sperm reserves was due to variation in male investment during mating.

We found that variation in the number of companion flies did not influence latency (Main effect of Conspecific number: *F_2,8_* = 0.249, *p*=0.785) (Figure 8A) or duration (Main effect of Conspecific number: *F_2,8_* = 0.14, *p*=0.873) of the first mating (Figure 8B). When females were allowed to remate, latency to mate was found to lengthen (Main effect of Mating number on mating latency: *F_1,6_* = 109.84, *p*=4.4 x 10^-5^) (Figure 8C) but duration of mating was unaffected (Main effect of Mating number on mating duration: *F_1,6_* = 4.001, *p*=0.092) (Figure 8D). However, neither was affected by the pattern of conspecific exposure (Effect of Mating number x Treatment on mating latency: *F_2,6_* = 3.16, *p*=0.116; Effect of Mating number x Treatment on mating duration: *F_2,6_* = 0.211, *p*=0.816).

**Figure 8.**
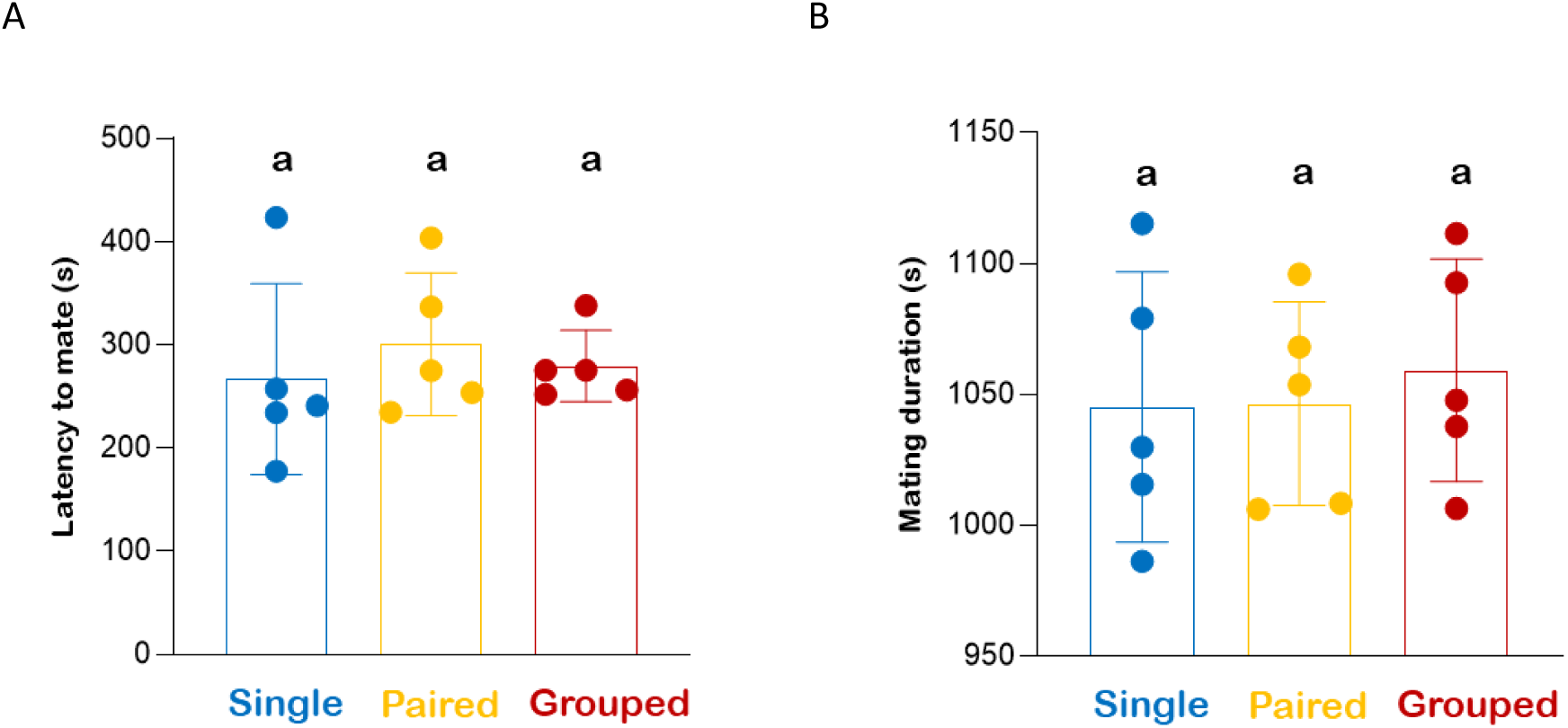
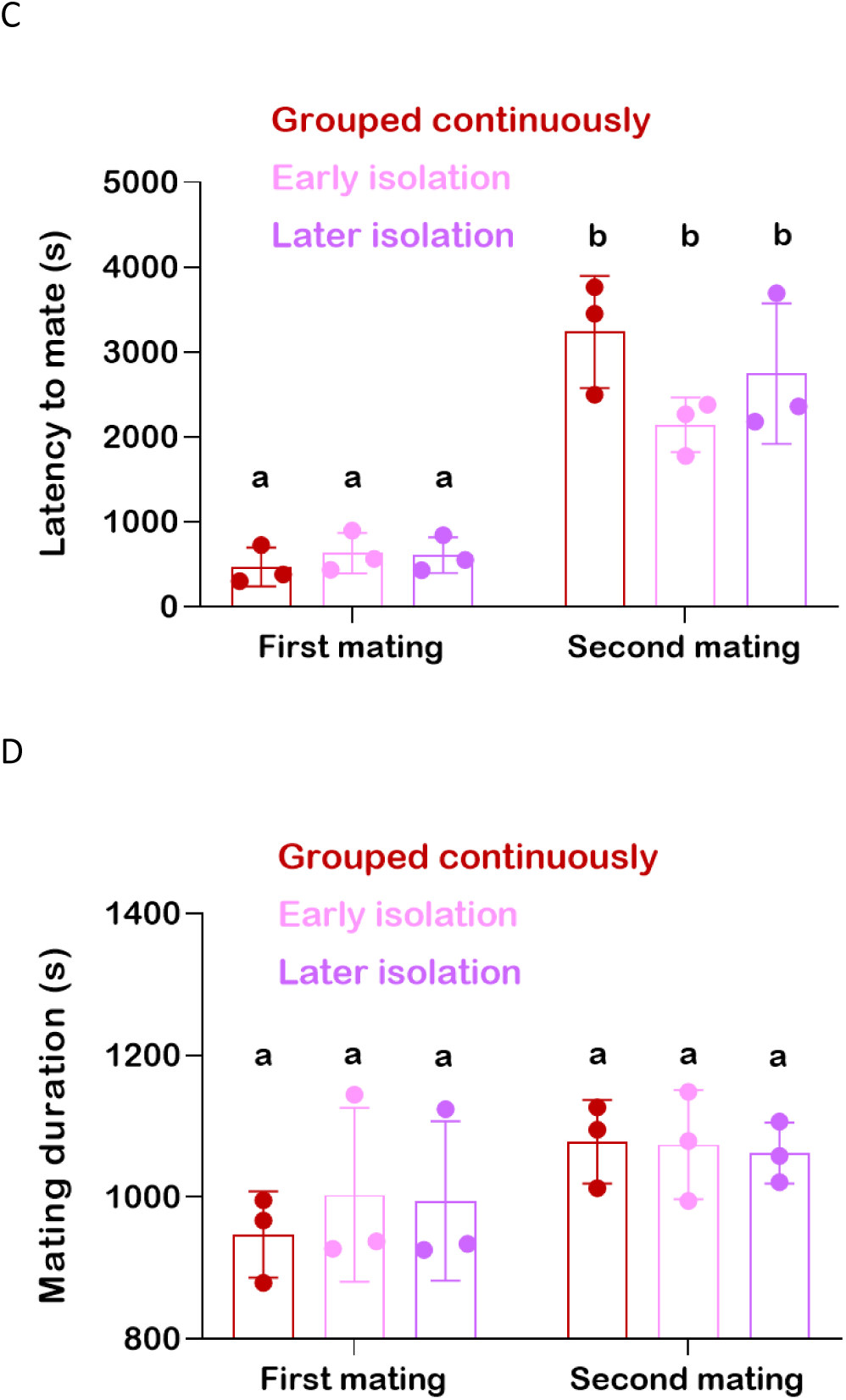
Mating latency and duration are unaffected by housing density. Mean values for mating latency and duration for Experiment 2 (A-B) and Experiment 3 (C-D) do not show differences across housing densities. Latency refers to the time taken to start copulation since introduction of a male to a mating vial containing a female, while duration represents the total time taken to complete copulation. Scatter points represent mean values for N replicate experiments, each obtained by averaging data across *n* individuals within the replicate. Height of each bar reflects the mean across N replicate experiments while error bars indicate the corresponding standard deviation. Bars which share any letter are not significantly different from one another. For A and B, data were analysed using a mixed-model ANOVA with Housing density as a fixed factor and Replicate experiment as random factor (N = 5 and *n* = 12 - 30 per treatment). For C and D, data were analysed using a repeated measures ANOVA with Housing density as a fixed factor and Mating number as a repeated measures factor (N = 3 and *n* varied between 5 - 9 per treatment). Pairwise comparisons in both experiments were performed using post-hoc Tukey’s HSD tests.

These results suggest that variation in progeny output cannot be attributed to variation in male reproductive investment during mating.

## 4. Discussion

### 4.1. Mechanisms underlying Allee effects of adult density in the fly

Negative effects of density are typically thought to result from competition among individuals for access to feeding and oviposition sites. Such effects can also result from inter-individual interactions between males and females which result in mate harm for the females (Linder and Rice, 2005; Orteiza et al., 2005; Edward et al., 2011; Fowler and Partridge, 1989). By contrast, mechanisms leading to positive density effects are less explored. The predominant hypothesis, inspired by studies on the flour beetle, *Tribolium confusum* (Park, 1933), attributes such effects to increasing frequency of matings with increases in density (Fujita, 1954; Watt, 1960; Rao et al., 2025). Since mating boosts fecundity in both *Tribolium* (Park, 1933) and *Drosophila* (Hanson and Ferris, 1929; Chen et al., 1988; Herndon & Wolfner, 1995), such increases in mating frequency are expected to increase reproductive output. However, this hypothesis fails to explain increased progeny output in our study as we controlled the number of matings across all treatments. Thus, while increased mating frequencies may contribute, they are certainly not necessary to observe Allee effects in the fly.

Since females vary their egg output in response to the presence of social cues around them (Churchill et al., 2021; Fowler et al., 2022), we considered that such plasticity may underlie the observed variation in lifetime reproduction across densities. We found evidence of reproductive plasticity in our experiments as females were able to increase or decrease progeny output depending on their immediate conspecific density. However, such plastic effects did not necessarily extend over longer periods of time. While previously grouped flies had diminished reproduction over their lifetimes after being isolated or paired, previously single or pair-housed flies showed only minor, short-lived increases in reproductive output upon grouping. We speculate that while these flies were able to react to density changes, they may be physiologically constrained in their ability to increase reproductive output. Such physiological barriers appeared to be lifted after remating, indicating that mating influences this density-dependent physiological state. It is unlikely that males exert much control over these mating dependent effects as the duration of mating, a commonly used proxy for male reproductive investment (Gilchrist and Partridge, 2000; Bretman et al., 2009), was comparable for both matings, and did not vary across densities. These effects must therefore be driven by density-dependent changes in the female herself.

We speculate that changes in storage of the male ejaculate by the female are a likely mechanism behind such effect. Single housing is known to reduce the levels of the neuropeptide *DH44* in fly brains (Zhao et al., 2024) which may lead to earlier sperm ejection and consequently reduced storage within the reproductive tract of transgenic females (Lee et al., 2015). Such reduced sperm reserves may explain lower progeny output in flies housed at low densities, as well as the trend for singly housed females to remate sooner than those housed in groups (Manning, 1962). Alternatively, other ejaculate components such as seminal fluid proteins may be stored differentially across densities. Components such as the sex peptide (SP) and accessory gland proteins (Acps) are known to modulate egg production by the female (Chen et al., 1988; Herndon & Wolfner, 1995). Interestingly, the effects of both SP and social context on oogenesis occur via modulation of juvenile hormone levels (Soller et al., 1999; Bailly et al., 2023), which may serve as a common physiological node for mating and density to influence egg output. A systematic assessment of sperm counts in the reproductive tracts of females as well as egg output across different densities is needed to compare the relative merits of these two hypotheses.

An important question to consider is *why* conspecific density might induce changes in storage of the ejaculate. Mated females attract other females to favourable oviposition sites by marking them with cVA, which is an aggregation pheromone (Duménil et al., 2016). Since cVA is produced only by males and acquired by females along with the male ejaculate (Butterworth, 1969; Bartelt et al., 1985), we speculate that females may sacrifice some of their stored ejaculate in the process of pheromone deposition. Given Allee effects associated with larval survival (discussed below), such communication is expected to increase local fly densities and consequently improve progeny survival. The benefits of such signalling are expected to be higher for females experiencing low densities who may deposit greater amounts of cVA, and consequently lose greater fractions of their stored sperm, at the oviposition site. In other words, females at low densities may trade-off future reproduction to secure the survival of their current progeny. Given that differences in pre-adult survival contributed little to variation in both total reproductive output as well as intrinsic growth rates, such behaviour appears to be non-adaptive. However, it is possible that under more natural ecologies, this trade-off is resolved more favourably.

In summary, our results suggest that repeated matings are not necessary for positive density effects on female reproduction to occur. Instead, these effects appear to be driven by density-dependent changes in the post-mating physiology of the female.

### 4.2. Effects of adult density on larval survival

Although positive relationships between larval density and pre-adult mortality are often observed under laboratory conditions (Sang, 1956; Gregg et al., 1990; Dombrovski et al., 2017; Rohlfs and Hoffmeister, 2004; Rohlfs et al. 2005; Trienens and Rohlfs, 2020), we found only weak support for such a relationship, as reductions in pre-adult mortality with increasing pre-adult densities were not statistically significant. Since such positive effects are thought to result from collective feeding interactions, the absence of such effects is not surprising as the food surface was scored with a scalpel to reduce the need for such interactions.

Interestingly, the density of adult females housed in a vial had significant effects on pre-adult mortality. Mortality was higher when females were paired, compared to when they were in groups indicating a positive effect. Similar effects were found by Wertheim et al. (2002), who suggested that these effects arose from suppression of fungal growth on the larval substrate at high adult densities. Such an explanation is unlikely to apply to our experiments, where propionic acid was added to the food media for inhibition of fungal growth (Ashburner and Roote, 2000). One explanation for such effects may be the presence of unfertilized eggs laid by virgin companion flies which may serve as an alternative source of nutrition (Ahmad et al., 2015). Additionally, adults may enhance the ability of larvae to feed by secreting amylases onto the food surface (Haj-Ahmad and Hickey, 1982). Such collective digestion is expected to be more efficient at higher densities leading to improved pre-adult survival at higher adult densities. Curiously, pre-adult mortality for progeny of singly housed females was lower than that seen for paired females, suggesting negative effects resulting from the presence of a single conspecific. While we are not aware of any previously described phenomena that may explain such a pattern, we speculate that singly housed flies may show plastic responses to the absence of conspecifics by investing more into their eggs. Larvae hatching from such eggs would have greater resources to burrow efficiently and may thus survive better. Further studies are needed to systematically test these hypotheses.

Broadly, our results support the idea that pre-adult survival is subject to Allee effects resulting from adult density. These may be of greater consequence than the effects of larval density, at least under some ecological conditions, and thus warrant detailed study.

### 4.3. Female reproduction is subject to Allee effects in Drosophila melanogaster

The canonical view of density effects in adult flies is one of negative effects on reproduction (Pearl and Parker, 1922; Robertson and Sang, 1944; Chiang and Hodson, 1950; Fujita 1954; Watt, 1960; Barker, 1973; Rodriguez 1989; Mueller and Huynh, 1994). Although positive density effects are known in other insect species (discussed in Watt, 1960), evidence in the fly is limited (Pearl, 1932; Roberson and Sang, 1944; Rockwell and Grossfield, 1978). Our results counter this view by providing strong, reproducible evidence that reproduction in female *Drosophila melanogaster* is positively affected by adult density. This idea is supported by recent work from Rao et al. (2025a, b) which reached the same conclusion using a different paradigm and different range of adult densities. Such variance across studies may be explained if density has both negative and positive effects on reproduction that are brought about by distinct mechanisms that vary across different experimental contexts.

For example, the overall level of nutrition is known to influence the magnitude of negative density effects in the fly (Robertson and Sang, 1944). Although most studies reporting negative density effects lacked yeast supplementation or employed low nutritive conditions for their assays, it is worth noting that nutrient supplementation does not guarantee expression of positive density effects (Robertson and Sang, 1944; Mueller and Huynh, 1994; Rao et al., 2025a). Spatial limitation on the food surface may be one explanation for the lack of positive effects under such conditions. Using a series of carefully designed experiments, Rao et al. (2025b) demonstrated that flies experienced negative density effects only in narrow sized vials while clear positive effects were evident in wider vials. However, this explanation fails to explain negative density effects in Robertson and Sang (1944), where the authors provided the experimental flies with an excess of yeast as well as large food surfaces.

We speculate that the continuous presence of males within the vial may be another factor that determines expression of positive density effects. Given that virtually all studies of density effects on reproduction in the fly have housed flies in mixed-sex groups with equal sex ratios, such effects are largely overlooked. To our knowledge, the only assessment of density effects on female reproductive output in the absence of males is by Rockwell and Grossfield (1978) which found evidence of positive density dependence, as seen in this study. Further studies that systematically vary the level of male presence experienced by females are needed to test if this correlation is causal in nature.

### 4.4. Allee effects in the fly: Proto-co-operation or ‘true’ co-operation?

A vast majority of the processes underlying Allee effects involve interactions among individuals. Density-dependent variation in the frequency or efficacy of co-operative interactions, such as collective defence or alloparental care, is an intuitive explanation for many instances of Allee effects (reviewed in Courchamp et al., 1999). However, as discussed above, Allee effects can also emerge from interactions such as mating, or indirect interactions such as niche modification (Allee and Bowen, 1932). These latter interactions may or may not benefit the actor but have incidental facilitatory effects on interactors. Such interactions are considered forms of ‘proto-cooperation’ (Allee et al., 1949; Courchamp et al., 1999), unlike ‘true’ cooperative interactions which have evolved to be beneficial to interactors, with or without benefits to the actor (Sachs et al., 2004; West et al., 2021). Despite their incidental nature, proto-co-operative interactions are of interest as they can represent early stages in the evolution of true co-operation via by-products (reviewed in Sachs et al., 2004).

While our results conclusively show the existence of Allee effects in *Drosophila melanogaster*, it remains to be seen whether the underlying mechanisms are facilitatory or co-operative in nature. As discussed previously, changes in the female’s physiology underlie these effects but it is uncertain what prompts these changes. One possibility is that absence of conspecifics at the oviposition site triggers physiological changes culminating in the deposition of cVA for signalling purposes. A loss of sperm in the process is an unintended consequence of the female’s inability to selectively deposit cVA. In this scenario, the presence of conspecifics simply facilitates lifetime reproductive output by eliminating the need for signalling. Another possibility is that these physiological changes are triggered by some form of active interaction between individuals. Female flies are known to use their wings to visually communicate information about parasitoid presence to other conspecifics which triggers physiological changes in the ovaries leading to suppression of egg laying (Kacsoh et al., 2015). Given the variety of inter-individual interactions known to occur in the fly, it is possible that females modulate their reproductive behaviour in response to social interactions. In this case, the magnitude of Allee effects is expected to depend on the levels of inter-individual interactions between flies, which may be tested by leveraging the vast genetic resources available in the fly (Anderson et al., 2016, 2017; Alwash et al., 2021). Presence of such a correlation would support the co-operative hypothesis while the absence would suggest the existence of facilitatory mechanisms.

Allee effects have been of interest to ecologists not merely because of their utility in understanding population dynamics, but also as an underlying driver of sociality (Allee et al., 1949; Stephens et al., 1999). Evidence of Allee effects in the fly suggest the possibility of adaptive benefits resulting from the diversity of inter-individual interactions that have been documented in the fly. Given the availability of numerous tools to track and measure behaviour in the fly (Branson et al., 2009; Kabra et al., 2013; Pereira et al., 2022), we expect that our understanding of behavioural interactions, their genetic underpinnings and ecological consequences will only increase with time. We, therefore, believe that the fly may prove to be a useful model to study how co-operation may emerge and shape social interactions.

## Acknowledgements

We thank Aarti Muralidharan for significant contributions to data collection as well as Sushma Rao, Navya M. and Vaibhav Jatolia for assistance during data collection. We also members of CBNL for critical feedback and Muniraju and Rajanna for laboratory support.

## References

Ahmad, M., Chaudhary, S. U., Afzal, A. J., & Tariq, M. (2015). Starvation-induced dietary behaviour in *Drosophila melanogaster* larvae and adults. Scientific Reports, 5, 14285.

Allee, W. C., & Bowen, E. S. (1932). Studies in animal aggregations: mass protection against colloidal silver among goldfishes. Journal of Experimental Zoology, 61(2), 185–207.

Allee, W. C., Emerson, A. E., Park, O., Park, T. & Schmidt, K. P. (1949). Principles of Animal Ecology. W. B. Saunders.

Alwash, N., Allen, A. M., Sokolowski, M. B., & Levine, J. D. (2021). The *Drosophila melanogaster* foraging gene affects social networks. Journal of Neurogenetics, 35(3), 249–261.

Anderson, B. B., Scott, A., & Dukas, R. (2016). Social behavior and activity are decoupled in larval and adult fruit flies. Behavioral Ecology, 27(3), 820–828.

Anderson, B. B., Scott, A., & Dukas, R. (2017). Indirect genetic effects on the sociability of several group members. Animal Behaviour, 123, 101–106.

Ashburner, M., Golic, K. G. & Hawley, R. S. (2005). Drosophila: A laboratory handbook (2nd ed.). Cold Spring Harbour Laboratory Press.

Bailly, T. P., Kohlmeier, P., Etienne, R. S., Wertheim, B., & Billeter, J. C. (2023). Social modulation of oogenesis and egg laying in *Drosophila melanogaster*. Current Biology, 33(14), 2865–2877.

Barker, J. S. F. (1973). Adult population density, fecundity and productivity in *Drosophila melanogaster* and *Drosophila simulans*. Oecologia, 11(2), 83–92.

Bartelt, R. J., Schaner, A. M., & Jackson, L. L. (1985). cis-Vaccenyl acetate as an aggregation pheromone in *Drosophila melanogaster*. Journal of Chemical Ecology, 11(12), 1747–1756.

Branson, K., Robie, A. A., Bender, J., Perona, P., & Dickinson, M. H. (2009). High-throughput ethomics in large groups of *Drosophila*. Nature Methods, 6(6), 451–457.

Bretman, A., Fricke, C., & Chapman, T. (2009). Plastic responses of male *Drosophila melanogaster* to the level of sperm competition increase male reproductive fitness. Proceedings of the Royal Society B: Biological Sciences, 276(1662), 1705–1711.

Butterworth, F. M. (1969). Lipids of *Drosophila*: a newly detected lipid in the male. Science, 163(3873), 1356–1357.

Chen, P. S., Stumm-Zollinger, E., Aigaki, T., Balmer, J., Bienz, M., & Böhlen, P. (1988). A male accessory gland peptide that regulates reproductive behavior of female *D. melanogaster*. Cell, 54(3), 291–298.

Chiang, H. C., & Hodson, A. C. (1950). An analytical study of population growth in *Drosophila melanogaster*. Ecological Monographs, 20(3), 173–206.

Churchill, E. R., Dytham, C., Bridle, J. R., & Thom, M. D. (2021). Social and physical environment independently affect oviposition decisions in *Drosophila*. Behavioral Ecology, 32(6), 1391–1399.

Courchamp, F., Clutton-Brock, T., & Grenfell, B. (1999). Inverse density dependence and the Allee effect. Trends in Ecology & Evolution, 14(10), 405–410.

del Solar, E., & Palomino, H. (1966). Choice of oviposition in *Drosophila melanogaster*. The American Naturalist, 100(911), 127–133.

Dombrovski, M., Poussard, L., Moalem, K., Kmecova, L., Hogan, N., Schott, E., Vaccari, A., Acton, S., & Condron, B. (2017). Cooperative behaviour emerges among *Drosophila* larvae. Current Biology, 27(18), 2821–2826.

Dukas, R. (2020). Natural history of social and sexual behavior in fruit flies. Scientific Reports, 10(1), 21932.

Duménil, C., Woud, D., Pinto, F., Alkema, J. T., Jansen, I., Van Der Geest, A. M., Roessingh. S., & Billeter, J. C. (2016). Pheromonal cues deposited by mated females convey social information about egg-laying sites in *Drosophila melanogaster*. Journal of Chemical Ecology, 42(3), 259–269.

Edward, D. A., Fricke, C., Gerrard, D. T., & Chapman, T. (2011). Quantifying the life-history response to increased male exposure in female *Drosophila melanogaster*. Evolution, 65(2), 564–573.

Emlen, D. J. (2008). The roles of genes and the environment in the expression and evolution of alternative tactics. In R. Oliveira, M. Taborsky & H. J. Brockmann (Eds.), Alternative Reproductive Tactics: An Integrative Approach. Cambridge University Press.

Ferreira, C. H., & Moita, M. A. (2020). Behavioral and neuronal underpinnings of safety in numbers in fruit flies. Nature Communications, 11(1), 4182.

Foley, B. R., Saltz, J. B., Nuzhdin, S. V., & Marjoram, P. (2015). A Bayesian approach to social structure uncovers cryptic regulation of group dynamics in *Drosophila melanogaster*. The American Naturalist, 185(6), 797–808.

Fowler, E. K., Leigh, S., Rostant, W. G., Thomas, A., Bretman, A., & Chapman, T. (2022). Memory of social experience affects female fecundity via perception of fly deposits. BMC Biology, 20(1), 244.

Fowler, K., & Partridge, L. (1989). A cost of mating in female fruitflies. Nature, 338(6218), 760–761.

Fujita, H. (1954). An interpretation of the changes in type of the population density effect upon the oviposition rate. Ecology, 35(2), 253–257.

Gilchrist, A. S., & Partridge, L. (2000). Why it is difficult to model sperm displacement in *Drosophila melanogaster*: The relation between sperm transfer and copulation duration. Evolution, 54(2), 534–542.

Gogna, N., Singh, V. J., Sheeba, V., & Dorai, K. (2015). NMR-based investigation of the *Drosophila melanogaster* metabolome under the influence of daily cycles of light and temperature. Molecular BioSystems, 11(12), 3305–3315.

Gregg, T. G., McCrate, A., Reveal, G., Hall, S., & Rypstra, A. L. (1990). Insectivory and social digestion in *Drosophila*. Biochemical Genetics, 28(3), 197–207.

Haj-Ahmad, Y., & Hickey, D. A. (1982). A molecular explanation of frequency-dependent selection in *Drosophila*. Nature, 299(5881), 350–352.

Hanson, F. B., & Ferris, F. R. (1929). A quantitative study of fecundity in *Drosophila melanogaster*. Journal of Experimental Zoology, 54(3), 485–506.

Herndon, L. A., & Wolfner, M. F. (1995). A *Drosophila* seminal fluid protein, Acp26Aa, stimulates egg laying in females for 1 day after mating. Proceedings of the National Academy of Sciences, 92(22), 10114–10118.

Hoffmeister, T. S., & Rohlfs, M. (2001). Aggregative egg distributions may promote species co-existence - but why do they exist? Evolutionary Ecology Research, 3(1), 37–50.

Iliadi, K. G., Iliadi, N. N., & Boulianne, G. L. (2009). Regulation of *Drosophila* life-span: effect of genetic background, sex, mating and social status. Experimental Gerontology, 44(8), 546–553.

Kabra, M., Robie, A. A., Rivera-Alba, M., Branson, S., & Branson, K. (2013). JAABA: Interactive machine learning for automatic annotation of animal behavior. Nature Methods, 10(1), 64–67.

Kacsoh, B. Z., Bozler, J., Ramaswami, M., & Bosco, G. (2015). Social communication of predator-induced changes in *Drosophila* behavior and germ line physiology. eLife, 4, e07423.

Lee, K. M., Daubnerová, I., Isaac, R. E., Zhang, C., Choi, S., Chung, J., & Kim, Y. J. (2015). A neuronal pathway that controls sperm ejection and storage in female *Drosophila*. Current Biology, 25(6), 790–797.

Leech, T., Sait, S. M., & Bretman, A. (2017). Sex-specific effects of social isolation on ageing in *Drosophila melanogaster*. Journal of Insect Physiology, 102, 12–17.

Leslie, P. H. (1945). On the use of matrices in certain population mathematics. Biometrika, 33(3), 183–212.

Linder, J. E., & Rice, W. R. (2005). Natural selection and genetic variation for female resistance to harm from males. Journal of Evolutionary Biology, 18(3), 568–575.

Manning, A. (1962). A sperm factor affecting the receptivity of *Drosophila melanogaster* females. Nature, 194(4825), 252–253.

Marks, R. W., Seager, R. D., & Barr, L. G. (1988). Local ecology and multiple mating in a natural population of *Drosophila melanogaster*. The American Naturalist, 131(6), 918–923.

McDonald, J.H. (2014). Handbook of Biological Statistics (3rd ed.). Sparky House Publishing.

Mery, F., Varela, S. A. M., Danchin, É., Blanchet, S., Parejo, D., Coolen, I., & Wagner, R. H. (2009). Public versus personal information for mate copying in an invertebrate. Current Biology, 19(9), 730–734.

Mueller, L. D., & Huynh, P. T. (1994). Ecological determinants of stability in model populations. Ecology, 75(2), 430–437.

Navarro, J., & del Solar, E. (1975). Pattern of spatial distribution in *Drosophila melanogaster*. Behavior Genetics, 5(1), 9–16.

Ochando, M. D., Reyes, A., & Ayala, F. J. (1996). Multiple paternity in two natural populations (orchard and vineyard) of *Drosophila*. Proceedings of the National Academy of Sciences, 93(21), 11769–11773.

Orteiza, N., Linder, J. E., & Rice, W. R. (2005). Sexy sons from re-mating do not recoup the direct costs of harmful male interactions in the *Drosophila melanogaster* laboratory model system. Journal of Evolutionary Biology, 18(5), 1315–1323.

Park, T. (1933). Studies in population physiology. II. Factors regulating initial growth of *Tribolium confusum* populations. Journal of Experimental Zoology, 65(1), 17–42.

Pearl, R. (1932). The influence of density of population upon egg production in *Drosophila melanogaster*. Journal of Experimental Zoology, 63(1), 57–84.

Pearl, R., & Parker, S. L. (1921). Experimental studies on the duration of life. I. Introductory discussion of the duration of life in Drosophila. The American Naturalist, 55(641), 481–509.

Pearl, R., & Parker, S. L. (1922). On the influence of density of population upon the rate of reproduction in *Drosophila*. Proceedings of the National Academy of Sciences, 8(7), 212–219.

Pearl, R., Miner, J. R., & Parker, S. L. (1927). Experimental studies on the duration of life. XI. Density of population and life duration in Drosophila. The American Naturalist, 61(675), 289–318.

Pereira, T. D., Tabris, N., Matsliah, A., Turner, D. M., Li, J., Ravindranath, S., Papadoyannis, E. S., Normand, E., Deutsch, D. S., Wang, Z. Y., McKenzie-Smith, G. C., Mitelut, C. C., Castro, M. D., D’Uva, J., Kislin, M., Sanes, D.H., Kocher, S. D., Wang, S. S. H., Falkner, A. L., Shaevitz, J. W., & Murthy, M. (2022). SLEAP: A deep learning system for multi-animal pose tracking. Nature methods, 19(4), 486–495.

Ramdya, P., Lichocki, P., Cruchet, S., Frisch, L., Tse, W., Floreano, D., & Benton, R. (2015). Mechanosensory interactions drive collective behaviour in *Drosophila*. Nature, 519(7542), 233–236.

Rao, M., Temura, C., Bindya, R. S., & Joshi, A. (2025b). Fitness effects of adult crowding in *Drosophila*: More than just overall density. bioRxiv, 10.1101/2025.04.24.650416.

Rao, M., Temura, C., Mital, A., Anvitha, S., & Joshi, A. (2025a). Bigger is not always better: Size-dependent fitness effects of adult crowding in *Drosophila melanogaster*. bioRxiv, 10.1101/2025.04.21.649761.

Robertson, F. W., & Sang, J. H. (1944). The ecological determinants of population growth in a *Drosophila* culture. I. Fecundity of adult flies. Proceedings of the Royal Society of London. Series B: Biological Sciences, 132(868), 258–277.

Rockwell, R. F., & Grossfield, J. (1978). *Drosophila*: Behavioral cues for oviposition. American Midland Naturalist, 99(2), 361–368.

Rodriguez, D. J. (1989). A model of population dynamics for the fruit fly *Drosophila melanogaster* with density dependence in more than one life stage and delayed density effects. The Journal of Animal Ecology, 58(2), 349–365.

Rohlfs, M., & Hoffmeister, T. S. (2004). Spatial aggregation across ephemeral resource patches in insect communities: an adaptive response to natural enemies? Oecologia, 140(4), 654–661.

Rohlfs, M., Obmann, B., & Petersen, R. (2005). Competition with filamentous fungi and its implication for a gregarious lifestyle in insects living on ephemeral resources. Ecological Entomology, 30(5), 556–563.

Ruan, H., & Wu, C. F. (2008). Social interaction-mediated lifespan extension of *Drosophila* Cu/Zn superoxide dismutase mutants. Proceedings of the National Academy of Sciences, 105(21), 7506–7510.

Sachs, J. L., Mueller, U. G., Wilcox, T. P., & Bull, J. J. (2004). The evolution of cooperation. The Quarterly Review of Biology, 79(2), 135–160.

Sang, J. H. (1956). The quantitative nutritional requirements of *Drosophila melanogaster*. Journal of Experimental Biology, 33(1), 45–72.

Sarin, S., & Dukas, R. (2009). Social learning about egg-laying substrates in fruitflies. Proceedings of the Royal Society B: Biological Sciences, 276(1677), 4323–4328.

Schneider, J., Dickinson, M. H., & Levine, J. D. (2012). Social structures depend on innate determinants and chemosensory processing in *Drosophila*. Proceedings of the National Academy of Sciences, 109(Supplement 2), 17174–17179.

Shultzaberger, R. K., Johnson, S. J., Wagner, J., Ha, K., Markow, T. A., & Greenspan, R. J. (2019). Conservation of the behavioral and transcriptional response to social experience among *Drosophilids*. *Genes*, Brain and Behavior, 18(1), e12487.

Simon, A. F., Chou, M.-T., Salazar, E. D., Nicholson, T., Saini, N., Metchev, S., & Krantz, D. E. (2012). A simple assay to study social behavior in *Drosophila*: measurement of social space within a group. *Genes*, Brain and Behavior, 11(2), 243–252.

Soller, M., Bownes, M., & Kubli, E. (1999). Control of oocyte maturation in sexually mature *Drosophila* females. Developmental Biology, 208(2), 337–351.

Stamps, J., Buechner, M., Alexander, K., Davis, J., & Zuniga, N. (2005). Genotypic differences in space use and movement patterns in *Drosophila melanogaster*. Animal Behaviour, 70(3), 609–618.

Stephens, P. A., & Sutherland, W. J. (1999). Consequences of the Allee effect for behaviour, ecology and conservation. Trends in Ecology & Evolution, 14(10), 401–405.

Stephens, P. A., Sutherland, W. J., & Freckleton, R. P. (1999). What is the Allee effect? Oikos, 87(1), 185–190.

Tinette, S., Zhang, L., & Robichon, A. (2004). Cooperation between *Drosophila* flies in searching behavior. *Genes*, Brain and Behavior, 3(1), 39–50.

Travis, J., Bassar, R. D., Coulson, T., Reznick, D., & Walsh, M. (2023). Density-dependent selection. Annual Review of Ecology, Evolution, and Systematics, 54(1), 85–105.

Trienens, M., & Rohlfs, M. (2020). A potential collective defense of *Drosophila* larvae against the invasion of a harmful fungus. Frontiers in Ecology and Evolution, 8, Article 79.

Verschut, T. A., Ng, R., Doubovetzky, N. P., Le Calvez, G., Sneep, J. L., Minnaard, A. J., Su, C., Carlsson, M. A., Wertheim, B., & Billeter, J. C. (2023). Aggregation pheromones have a non-linear effect on oviposition behavior in *Drosophila melanogaster*. Nature Communications, 14(1), 1544.

Watt, K. E. (1960). The effect of population density on fecundity in insects. The Canadian Entomologist, 92(9), 674–695.

Wertheim, B., Allemand, R., Vet, L. E., & Dicke, M. (2006). Effects of aggregation pheromone on individual behaviour and food web interactions: a field study on *Drosophila*. Ecological Entomology, 31(3), 216–226.

Wertheim, B., Dicke, M., & Vet, L. E. (2002). Behavioural plasticity in support of a benefit for aggregation pheromone use in *Drosophila melanogaster*. Entomologia Experimentalis et Applicata, 103(1), 61–71.

West, S. A., Cooper, G. A., Ghoul, M. B., & Griffin, A. S. (2021). Ten recent insights for our understanding of cooperation. Nature Ecology & Evolution, 5(4), 419–430.

Zhao, H., Jiang, X., Ma, M., Xing, L., Ji, X., & Pan, Y. (2024). A neural pathway for social modulation of spontaneous locomotor activity (SoMo-SLA) in *Drosophila*. Proceedings of the National Academy of Sciences, 121(9), e2314393121.

