## Supplementary for "Density-dependent plasticity of female reproductive output mediates Allee effects in *Drosophila melanogaster*"

**Supplementary information**

**Supplementary methods**

**<u>1. Construction of Leslie matrices</u>**

The life history of females over the course of the experiment is described again with the day of egg laying as day 0. Emergence occurs on day 9 while reproduction begins after mating on day 12. Experimental flies are transferred to new vials every day for the first 7 days post mating and every two days thereafter. For assays where the mated fly was tracked, mortality was checked along with each transfer at the start of transfer period. All egg laying occurred after this transfer. Thus, this represents a pre-reproductive census scheme where time units change just before the birth pulse.

The Leslie matrix is a square matrix (i.e. number of rows and columns are equal). Each column stores information about a specific age class (day in this case). Since a pre-reproductive census is employed, the first row of the matrix represents net fecundity values i.e. products of fecundity and the hatching rate for each age. The rest of the matrix consists of zeros except the lower sub diagonal which stores fraction of individuals in a given age class surviving to the next age class.

$$M = \begin{bmatrix} F_0 & F_1 & F_2 & \cdots & \cdot & F_k & F_{k+1} & \cdots & F_{m-1} & F_m \\ P_0 & \cdot & \cdot & \cdots & \cdot & \cdot & \cdot & \cdots & \cdot & \cdot \\ \cdot & P_1 & \cdot & \cdots & \cdot & \cdot & \cdot & \cdots & \cdot & \cdot \\ \cdot & \cdot & P_2 & \cdots & \cdot & \cdot & \cdot & \cdots & \cdot & \cdot \\ \cdots & \cdots & \cdots & \cdots & \cdots & \cdots & \cdots & \cdots & \cdots & \cdots \\ \cdot & \cdot & \cdot & \cdots & P_{k-1} & \cdot & \cdot & \cdots & \cdot & \cdot \\ \cdot & \cdot & \cdot & \cdots & \cdot & P_k & \cdot & \cdots & \cdot & \cdot \\ \cdots & \cdots & \cdots & \cdots & \cdots & \cdots & \cdots & \cdots & \cdots & \cdots \\ \cdot & \cdot & \cdot & \cdots & \cdot & \cdot & \cdot & \cdots & P_{m-1} & \cdot \end{bmatrix} \quad 0 < P_x < 1;\ F_x \geqq 0.$$

(Modified from Leslie, 1945)

This matrix can be trimmed to exclude the columns beyond the age of reproduction (k) to obtain a matrix A that is relevant to understanding growth of the population.

$$A = \begin{bmatrix} F_0 & F_1 & F_2 & F_3 & \cdots & F_{k-1} & F_k \\ P_0 & \cdot & \cdot & \cdot & \cdots & \cdot & \cdot \\ \cdot & P_1 & \cdot & \cdot & \cdots & \cdot & \cdot \\ \cdot & \cdot & P_2 & \cdot & \cdots & \cdot & \cdot \\ \cdots & \cdots & \cdots & \cdots & \cdots & \cdots & \cdots \\ \cdot & \cdot & \cdot & \cdot & \cdots & P_{k-1} & \cdot \end{bmatrix}.$$

(Modified from Leslie, 1945)

Leslie (1945) showed that this matrix can be transformed to a different coordinate space using any transformation matrix H to obtain a new matrix B that is equivalent to A but operates in the new frame of reference. When H has the following form,

$$H = \begin{bmatrix} (P_0 P_1 P_2 \dots P_{k-1}) & \cdot & \cdot & \dots & \cdot & \cdot & \cdot \\ \cdot & (P_1 P_2 P_3 \dots P_{k-1}) & \cdot & \dots & \cdot & \cdot & \cdot \\ \cdot & \cdot & (P_2 P_3 \dots P_{k-1}) & \dots & \cdot & \cdot & \cdot \\ \dots & \dots & \dots & \dots & \dots & \dots & \dots \\ \cdot & \cdot & \cdot & \dots & (P_{k-2} P_{k-1}) & \cdot & \cdot \\ \cdot & \cdot & \cdot & \dots & \cdot & P_{k-1} & \cdot \\ \cdot & \cdot & \cdot & \dots & \cdot & \cdot & 1 \end{bmatrix}$$

(Modified from Leslie, 1945)

the matrix A takes the following form

$$B = HAH^{-1} = \begin{bmatrix} F_0 & P_0 F_1 & P_0 P_1 F_2 & P_0 P_1 P_2 F_3 & \dots & (P_0 P_1 P_2 \dots P_{k-1}) F_k \\ 1 & \cdot & \cdot & \cdot & \dots & \cdot \\ \cdot & 1 & \cdot & \cdot & \dots & \cdot \\ \cdot & \cdot & 1 & \cdot & \dots & \cdot \\ \cdot & \cdot & \cdot & 1 & \dots & \cdot \\ \dots & \dots & \dots & \dots & \dots & \dots \\ \cdot & \cdot & \cdot & \cdot & \dots 1 & \cdot \end{bmatrix}.$$

(Modified from Leslie, 1945)

where all survivorship values are replaced with unity and the first row consists of the net fecundity values for different ages multiplied by the probability of surviving until the corresponding age, i.e. a product of all age specific survival fractions until that age ($P_0$*$P_1$*…$P_{x-1}$). Although the population represented by this transformed matrix is different from original one (for eg. in this case, all individuals survive until the last age of reproduction but die en masse after that), the latent roots of both matrices remain the same such that the intrinsic growth rate (λ) is unchanged.

For the Experiment 1, no information is available for adult mortality and only cumulative mortality over the pre-adult stages is known (from estimates of pre-adult survival). Fecundity estimates are available for all ages, but these are biased towards zero as dead flies are not excluded before averaging (and hence have zero fecundity). These measurements are insufficient for measurement of λ using the conventional form of the Leslie matrix but can be employed successfully using the transformed matrix B described above (along with a simplifying assumption).

The first row of the B form of the Leslie matrix discussed above consists of values of the form – $P_0$*$P_1$*…$P_{x-1}$ * $F_x$ where $P_x$ represents the probability of survival to the x+1 time point of all individuals that are alive at the x timepoint and $F_x$ represents the mean number of progeny produced by a female at timepoint x that hatch successfully (i.e. enter the first age class). Since $F_x$ is zero for all pre-adult stages, the first-row terms for all pre-adult stages can be set to 0. For adult stages, the above expression can be rewritten as $P_a$* $P_X$*$F_x$ where $P_a$ is the probability of reaching adulthood and $P_X$ is the probability of an adult surviving until age x (and is the product of all ages after reaching adulthood until age x-1 i.e. $P_{10}$*…$P_{x-1}$). To obtain $P_X$ from the observed data, we need the ratio of flies that survived to age x ($n_x$) to the total number of flies (N) assayed. To obtain $F_x$ from the observed data, we need to measure the ratio of total progeny produced at age x that hatched successfully (hatching rate ($h_x$) x number of eggs laid ($e_x$)) to the total number of flies alive at age x ($n_x$). Thus, the first row terms take the form –

$P_a * P_X * F_x = P_a * (n_x / N) *( h_x * e_x / n_x) = P_a * h_x * e_x /N$ – equation 1

If we make a simplifying (but possibly incorrect) assumption that the total number of progeny observed in a vial (dead or alive) represents the total number of eggs laid, then $e_x$ becomes $(a_x + d_x)$ while $P_a * h_x$ represents the fraction of eggs that survive to adulthood i.e. $(a_x / (a_x + d_x))$ and the equation 1 reduces to-

$$P_a * P_X * F_x = a_x / (a_x + d_x) * (a_x + d_x) /N$$

$$= a_x /N$$

which represents the mean number of progeny that emerge successfully.

Given these data are available for each density treatment, separate lifetables could be constructed for each treatment and the corresponding λ estimated.

**2. Randomization tests to compare intrinsic growth rates across treatments**

To test if the estimated λ values were significantly different across the three treatments, we employed randomization tests which assumed that the three datasets were obtained by random sampling from the same population. Under such a null hypothesis, individuals assigned to each treatment are exchangeable and can thus be shuffled across treatments without appreciably changing the estimates of λ. Thus, if the null hypothesis is true, the observed variation in λ values across treatments should be within the range of variation observed when such shuffled or randomized datasets are used. We constructed randomized datasets by randomly shuffling individuals across treatments and averaging progeny counts in each instance. These mean progeny counts were used to obtain estimates of λ under the null hypothesis. 1000 such randomizations were performed and observed variance in λ was compared to expected variance in λ across such randomizations. The null hypothesis was rejected if the observed value lay within the extreme 5% of the randomized values. A similar procedure was employed to make pairwise comparisons by using the difference between pairs of λ values as the test statistic.

**3. Randomization tests to assess contribution of different fitness components to variation in intrinsic growth rates across treatments**

As discussed above, the life table is constructed using information on progeny output across ages as described in equation 1 above. However, it can also be expressed as a product of three distinct components of fitness – pre-adult survival, distribution of reproductive effort across time and total reproductive output.

$P_a * P_X * F_x = P_a * h_x * e_x / N$

$= ((a_x / e_x) * (e_x * E/E))/N$ --------- $P_a * h_x = a_x / e_x$ and $E = \Sigma e_x$

$= (a_x / e_x) / N * (e_x / E)/N * E/N$

$(a_x / e_x) / N$ represents the mean survival to adulthood for eggs laid at age x, $(e_x / E)/N$ represents the mean fraction of reproductive effort at age x and E/N represents the mean lifetime reproductive output.

Since information is available about each of these components, an alternative way of constructing the first row is by multiplying these data. While this exercise yields the same mean progeny values, and consequently the same λ values, it also allows for randomization of individual fitness components. Such a randomization exercise can help parse the contribution of individual components of fitness to growth rates. For example, if pre-adult survival and lifetime reproductive output data are randomized across individuals without randomizing data for age-specific reproductive effort, then the randomized λ for a given treatment is attributable only to distribution of reproductive effort associated with that treatment. The observed λ can be compared to a distribution of 1000 such randomized values to determine the probability of obtaining the observed value solely due to treatment specific patterns of reproductive effort across time.

We assessed the contribution of different fitness components to λ using null hypotheses indicated in Supplementary Table 1. The first hypothesis tests if treatment-specific variation in any of the fitness components is essential to obtain the observed λ value for any given treatment. If rejected, then the subsequent hypotheses are tested to assess sufficiency and necessity of each fitness component for obtaining the observed λ. Hypotheses 2-4 assume sufficiency of individual fitness components for producing the observed λ. Thus, if the observed λ is found to be more extreme than randomized values, then the particular component of fitness can be considered insufficient. Hypotheses 5-7 assume that individual components of fitness are not necessary for producing the observed λ. Rejection of these hypotheses indicates that a given component of fitness is necessary to produce the observed λ.

Supplementary Table 1.

| Null hypothesis | | Randomization status under the null | | |
|---|---|---|---|---|
| | | Lifetime reproductive output | Distribution of reproductive effort | Pre-adult mortality survival |
| 1. | Variation in components of fitness is not necessary to produce the observed λ | + | + | + |
| 2. | Variation in lifetime reproductive output is sufficient to produce the observed λ | - | + | + |
| 3. | Variation in distribution of reproductive effort is sufficient to produce the observed λ | + | - | + |
| 4. | Variation in pre-adult survival is sufficient to produce the observed λ | + | + | - |
| 5. | Variation in lifetime reproductive output is not necessary to produce the observed λ | + | - | - |
| 6. | Variation in distribution of reproductive effort is not necessary to produce the observed λ | - | + | - |
| 7. | Variation in pre-adult survival is not necessary to produce the observed λ | - | - | + |

Table 1. Null hypotheses for testing sufficiency and necessity of contribution from individual components of fitness towards the observed growth rates.

Age-specific progeny counts are a function of lifetime reproduction, i.e. the total number of dead or live progeny produced, by a female, the distribution of reproductive effort across time, and maternal age-specific mortality experienced by progeny in the pre-adult stages. Distinct null hypotheses were proposed to assess the contribution of a component of fitness to the observed λ. Any components of fitness randomized under a given hypothesis are indicated by '+' while those left unaffected are indicated by '-'.

Supplementary figure 1

A

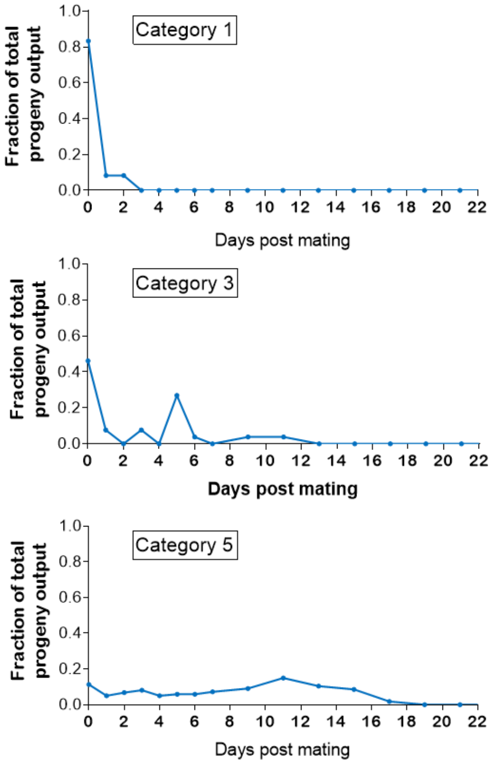


B

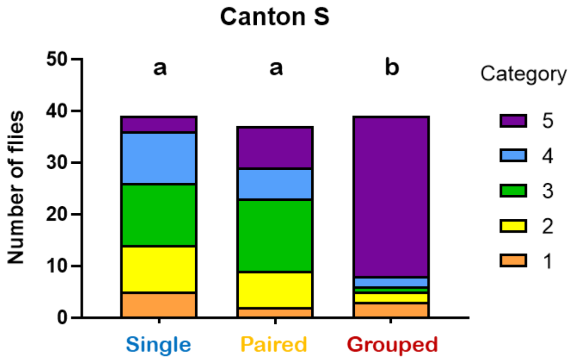


C

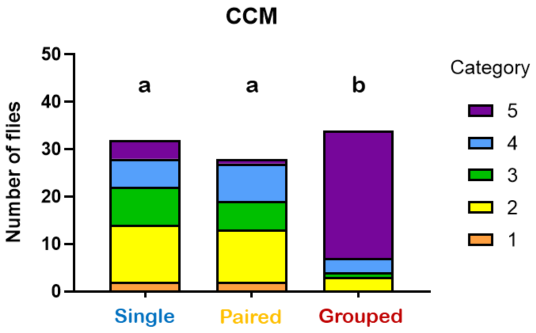

**Supplementary Figure 1. Individual patterns of progeny output across time change with housing density**

A) Normalized distributions of progeny output across time for individual females can be qualitatively classified into five shape categories. Category 1 exhibits a single peak that occurs shortly after mating after which no progeny emerge successfully. Category 2 shows an additional peak after this initial peak, which may occur at any age, while Category 3 exhibits multiple such secondary peaks. Distributions where such secondary peaks were less distinct and appear fused were classified into Category 4 while Category 5 represented individuals with largely continuous progeny output and limited consolidation across time.

Abundance of different types of progeny output distributions is significantly different for grouped flies compared to singly housed and paired flies in both (B) Canton S and (C) CCM backgrounds. Height of each bar reflects the total number of individuals per treatment, pooled across N replicate experiments while the height of each subdivided bar represents the number of flies in any given category. Bars which share any letter do not differ significantly in the distribution of categories. chi-square tests were used for omnibus testing and pairwise comparisons (N = 4 and 3 for Canton S and CCM respectively, *n* = 8 - 10 per treatment).

---

Supplementary Figure 2

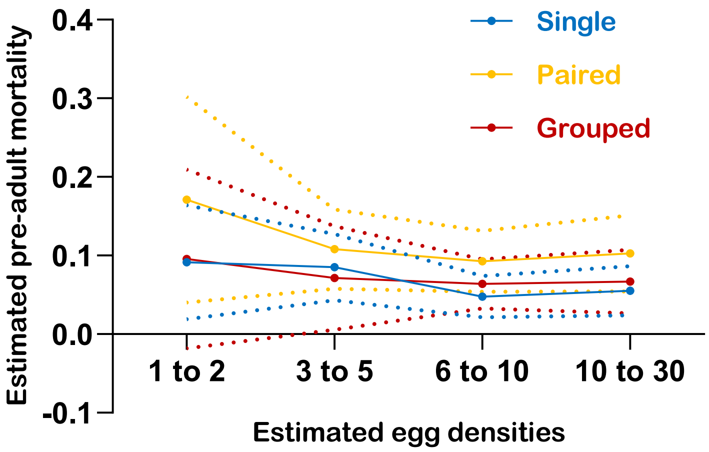


**Supplementary figure 2. Effect of pre-adult density on pre-adult mortality is not significant.**

Progeny mortality during the pre-adult does not vary significantly across estimated egg densities. Egg density was estimated as the total number of dead or live progeny found in a vial. Mortality rates across days and individuals were sorted according to their associated egg density and mean mortality for different egg densities was calculated. Each point on a line represents the mean value across replicate experiments and strains while dotted lines indicate standard deviation across N replicate experiments. Data analysed using a repeated measures ANOVA with Housing density and Strain as fixed factors and Egg density as the repeated measures factor (N = 4 and 3 for Canton S and CCM respectively, *n* = 8 - 10 per treatment).
